# Distinct within-host bacterial populations ensure function, colonization and transmission in leaf symbiosis

**DOI:** 10.1101/2021.12.06.471530

**Authors:** Tessa Acar, Sandra Moreau, Olivier Coen, Frédéric De Meyer, Olivier Leroux, Marine Beaumel, Paul Wilkin, Aurélien Carlier

**Affiliations:** Laboratory of Microbiology, Ghent University, 9000 Ghent, Belgium; LIPME, Université de Toulouse, INRAE, CNRS, 31320 Castanet-Tolosan, France; Ghent University, Department of Biology, K. L. Ledeganckstraat 35, 9000 Gent, Belgium; School of Built Environment and Bioeconomy, Tampere University of Applied Sciences, P.O. Box 356, 33101 Tampere, Finland; Royal Botanical Gardens Kew, Richmond, London, TW9 3AE, United Kingdom

## Abstract

Hereditary symbioses have the potential to drive transgenerational effects, yet the mechanisms responsible for transmission of heritable plant symbionts are still poorly understood. The leaf symbiosis between *Dioscorea sansibarensis* and the bacterium *Orrella dioscoreae* offers an appealing model system to study how heritable bacteria are transmitted to the next generation. Here, we demonstrate that inoculation of apical buds with a bacterial suspension is sufficient to colonize newly-formed leaves and propagules, and to ensure transmission to the next plant generation. Flagellar motility is not required for movement inside the plant, but is important for the colonization of new hosts. Further, stringent tissue-specific regulation of putative symbiotic functions highlight the presence of two distinct subpopulations of bacteria in the leaf gland and at the shoot meristem. We propose that bacteria in the leaf gland dedicate resources to symbiotic functions, while dividing bacteria in the shoot tip ensure successful colonization of meristematic tissue, glands and propagules. Compartmentalization of intra-host populations, together with tissue-specific regulation may serve as a robust mechanism for the maintenance of mutualism in leaf symbiosis.

**Importance:** Several plant species form associations with bacteria in their leaves, called leaf symbiosis. These associations are highly specific, but the mechanisms responsible for symbiont transmission are poorly understood. Using the association between the yam species *Dioscorea sansibarensis* and *Orrella dioscoreae* as a model leaf symbiosis, we provide experimental evidence that bacteria are transmitted vertically and distributed to specific leaf structures via association with shoot meristems. Flagellar motility is required for initial infection, but does not contribute to spread within host tissue. We also provide evidence that bacterial subpopulations at the meristem or in the symbiotic leaf gland differentially express key symbiotic genes. We argue that this separation of functional symbiont populations, coupled to tight control over bacterial infection and transmission, explain the evolutionary robustness of leaf symbiosis. These findings may provide insights into how plants may recruit and maintain beneficial symbionts at the leaf surface.

## Introduction

Heritable symbioses are common in animals, with many examples in invertebrates. For example, aphids (Hemiptera) harbor *Buchnera* bacteria, and 16% of all known insect species interact with *Wolbachia* bacteria (1–3). These model systems have provided tremendous insights into the cellular mechanisms underlying heritable symbiont transmission (4–6). In contrast to animal symbioses, most well-described plant-microbe symbioses rely on horizontally-transmitted symbionts, such as the interactions involving rhizobia or mycorrhizal fungi (7). Heritable transmission of symbionts has been demonstrated for only a handful of plant taxa, and the mechanisms governing symbiont transmission are still poorly understood (8, 9). However, recent evidence suggests that vertically-transmitted symbionts may also account for important transgenerational phenotypes (10–12). In addition, mode of transmission has important implications for the evolution of host-microbe associations. Indeed, while horizontal transmitted symbionts are usually vetted through a combination of partner choice and sanctions and rewards, vertical transmission is thought to be an efficient mechanism to establish successful cooperation through partner fidelity feedback (13, 14).

Three Angiosperm families include species that harbor possible vertically-transmitted bacterial symbionts (15). In the Primulaceae family, 30 out of 35 species of *Ardisia* display small glands at the leaf margin, colonized by *Burkholderia* bacteria (16). Congruent phylogenies of host and symbiotic bacteria suggest co-speciation and a strictly vertical mode of transmission (17). In the Rubiaceae family, nearly 500 plant species engage in leaf nodule symbiosis, with about 350 species in the *Pavetta* genus, 85 in *Psychotria* and 12 in *Sericanthe* (18–20). Unlike the Primulaceae family, the structures housing the bacteria have variable morphologies and may be distributed throughout the leaf lamina or along the midvein (19). Similar to *Ardisia*, the symbiosis involves specific associations with *Burkholderia* bacteria. Here, phylogenetic patterns suggest a mixed mode of transmission, with vertical transmission and occasional events of host-switching (16, 21, 22). In addition, the symbionts of *Psychotria punctata* are present in all life stages of the plant, including flower buds, anthers, gynoecium and embryos, providing strong evidence of vertical transmission in this taxon (23). Leaf nodule bacteria lack the genetic ability to fix nitrogen or metabolize phytohormones. Instead, the symbionts provide secondary metabolites that may protect the host against phytophagous insects or competitors (15, 24, 25). Because of their mutual dependence, the study of the molecular mechanisms underlying the associations between heritable leaf nodule bacteria and their hosts is challenging. For example, aposymbiotic seeds of *Psychotria* sp. and *Ardisia crenata* germinate normally, but fail to develop more than a few leaves and do not reach maturity (26). Moreover, genomes of leaf nodule bacteria do not encode known signalling pathways such as Nod factors, type III secreted effectors or plant hormones (15), and the molecular functions enabling colonization and transmission are unknown.

More than thirty years ago, Miller and Reporter used microscopy techniques and described the presence of a bacterial symbiont in the leaf acumen of *Dioscoreae sansibarensis*, but did not identify the bacterium (Miller & Reporter, 1987). Interestingly, symbiont-free plants could reportedly be obtained by surface-sterilization of bulbils, although these aposymbiotic plants readily became colonized by bacteria upon transfer to a non-sterile environment. We recently isolated and described these symbiotic bacteria as *Orrella dioscoreae* (*Alcaligenaceae*) (28, 29). Leaves of *D. sansibarensis* are heart-shaped and end with a distal acumen or forerunner-tip which exclusively harbours *O. dioscoreae*. In contrast to symbionts of Rubiaceae and Primulaceae, *O. dioscoreae* can be cultured outside of the host plant and is amenable to genetic manipulation (29). *O. dioscoreae* can be isolated from vegetative propagules and recent data indicates that the association with *D. sansibarensis* is ubiquitous throughout the range of the host plant. This suggests a vertical mode of transmission, but low phylogenetic congruence between plant and symbiont genetic markers indicates possible horizontal or host-switching transmission (30). Moreover, genomes of *O. dioscoreae* strains do not display any of the hallmarks of genome reductive evolution, a common phenomenon in vertically transmitted leaf symbioses (15).

In this work, we show that the association between *D. sansibarensis* and *O. dioscoreae* is tissue-specific. Using a newly developed gnotobiotic system, we demonstrate that bacteria are transmitted vertically, with possible horizontal transmission relying on bacterial motility and infection of developing apical buds. Our results provide insights into the transmission of a heritable bacterial symbiont in land plants and some of the molecular mechanisms that shape the evolution of leaf-bacteria symbioses.

## Material and Methods

### Plant culture and propagation

Plants were maintained in the greenhouse of the Laboratory of interactions Plant-Microbe-Environment (LIPME) in Castanet-Tolosan, France. Unless otherwise indicated plants were grown in climate chambers at 28°C, 70% humidity and a light cycle of 16h light (210 μmol/m^2^/s), 8h dark. Chemicals and reagents were purchase from Merck, France unless otherwise indicated. Micropropagation of *Dioscorea sansibarensis* was done using a protocol adapted from Alizedah et al. (31). Node cuttings were collected from greenhouse-grown plants after 2-4 months of growth.

Explants were surface sterilized by submerging them in a 5% solution of Plant Preservative Mixture (PPM, Plant Cell Technology, USA) with shaking at 100 rpm for 8hrs at 28°C, in the dark. After 8 hours, the bleached extremities of the explants were removed with a sterile scalpel. Explants were placed in sterilized growth medium (Murashige and Skoog basal salts (MS): 4.4g/L, 2% sucrose, vitamins: glycine (2mg/L), myo-inositol (100 mg/L), nicotinic acid (0.5mg/L), pyridoxine-HCl (0.5mg/L), thiamine.Cl (0.1mg/L) and L-cystein (20mg/L), pH=5.7), supplemented with 200 μg/ml carbenicillin (Meridis, France), 200 μg/ml cefotaxime (Meridis, France) and 0.2% v/v plant preservative mixture (PPM, Plant Cell Technology, USA). Explants were incubated at 28°C, 16h/8h of light cycle for 10 days. The medium was refreshed after 10 days, including supplements and antibiotics. After 21 days of incubation, the medium was replaced with growth medium containing MS, sucrose, PPM and vitamins as described above but without the antibiotics. Cuttings were transferred in magenta GA-7 vessels (Merck), incubated at 28°C, 16h of light until rooting.

### Detection and identification of bacteria

The tip of the leaf was dissected with tweezers and a scalpel, and the tissue was homogenized using 100 μl 0.4% NaCl and 3 sterile glass beads for 1 minute at 30 Hz in a ball mill (Retsch MM 400). The homogenized suspension was centrifuged briefly to pellet debris. One hundred μL of supernatant was directly plated out on TSA (Sigma) plates and incubated for 2 days at 28°C. If the plate showed growth, one isolate per colony type was picked and identified using colony PCR with primers specific for *O. dioscoreae* (nrdA-01-F, nrdA-02-R, Table S2), or with universal 16S rRNA primers (pA and pH, Table S2) followed by Sanger sequencing.

### Inoculation of *D. sansibarensis* with bacteria

Node cuttings were grown in axenic conditions (25ml MS + 2% sucrose + 0.2% PPM in Magenta vessel, 28°C, 16h/8h light cycle) until a new shoot appeared (after 6 weeks approximately). Verified aposymbiotic plants (tested as stated above) were inoculated with a strain of interest as followed: bacterial cultures in the exponential phase of growth were centrifuged (5000 rpm, 10 min) and washed twice with sterile 0.4% NaCl. Cell suspensions were normalized to OD_600nm_ = 0.2. The biggest leaf at the apical bud was gently pushed aside and 2 μl of the bacterial suspension (OD_600nm_ = 0.2) was gently deposited onto the apical bud (Suppl Figure 1). Plants were transferred to sterile microboxes (50ml MS + 2% sucrose + 0.2% PPM) at 28°C, 16h of light until new leaves emerged. Colonization was evaluated by dissecting a leaf tip and spreading the contents on suitable microbiological medium as described above. Plants were transferred to pots with soil and incubated in growth chamber. Shortly before senescence, plants develop bulbils. These bulbils were harvested and stored in a dark, dry place at room temperature for about 6 months or until dormancy broke. Sprouting bulbils were planted in soil and pots were left at 25°C, 16h of light.

### Bacterial genetics

*O. dioscoreae* strain R-71412 is a spontaneous nalidixic acid-resistant strain derived from *O. dioscoreae* LMG 29303^T^ (29). To obtain strain R-71417, a mini-Tn7 cassette containing the mCherry reporter gene was introduced into *O. dioscoreae* R-71412 by tri-parental mating as in Choi and Schweizer (32). Briefly, overnight cultures of recipient (*O. dioscoreae* R-71412), donor (*E. coli* S17-1 mini-Tn7::*mCherry*) and helper strain (*E. coli* S17-1 pUX-BF13) were diluted 1:100 in fresh medium without antibiotics (LB for *E. coli* and TSB for *O. dioscoreae*) and grown to OD_600nm_ = 0.5 while shaking at 37°C or 28°C. Cells were washed once in sterile 0.9% sodium chloride and re-suspended in sterile LB medium to OD_600nm_ ~ 1. About 100 μL of each suspension was spotted on LB agar without antibiotics and incubated overnight at 37°C. Cells were suspended in 500 μl of 0.9% NaCl solution and plated on selective medium (TSA supplemented with nalidixic acid (30 μg/mL) and gentamycin (20 μg/mL)) and incubated at 30 °C for 48h. Fluorescent colonies were visualized with a stereomicroscope (Leica DFC 7000T). The insertion of the transposon downstream of the *glmS* gene was confirmed by PCR using primers “Mini Tn7 primer forward” and “Mini Tn7 primer reverse” (Table S2).

To create a motility impaired *O. dioscoreae* mutant, a mutant allele of a *motB* homolog (locus tag ODI_R2122) was created by PCR amplification of three overlapping DNA fragments, containing the flanking regions of the gene of interest and the kanamycin resistance cassette from pKD4 (33). The upstream flanking region of the *motB* gene was amplified by using primers motB-UpF-GW and motB-UpR-kan, and the downstream flanking region of the *motB* gene was amplified by using primers motB-DnF-kan and motB-DnR-GW (Table S2). The up- and downstream fragments were fused together and amplified by using primers GW-*attB*1 and GW-*attB*2 (Table S2) by overlap extension PCR (SOE PCR) to generate the *motB* mutant allele. Once the fragments were verified, the PCR constructs were ligated into pDONRPEX18Tp-SceI-pheS (34) using the Invitrogen BP ligation kit and transformed by electroporation into *E. coli* Top 10. Suicide plasmids were introduced in *O. dioscoreae* (R71417) by triparental mating as above, using *E. coli* harbouring plasmid pRK600 as helper. Transconjugants were selected on TSA medium supplemented with kanamycin 50 μg/ml and nalidixic acid 30 μg/ml and incubated for 2 days at 28°C. Counter-selection of merodiploid clones was done by spreading on AB minimal medium supplemented with 0.2% citrate, 0.1% yeast extract and 0.1% (wt/vol) *p*-chlorophenylalanine (cPhe) (DL-4-chlorophenylalanine; Sigma-Aldrich). Colonies were screened for loss of trimethoprim resistance on TSA medium supplemented with nalidixic acid 30 μg/ml and kanamycin 30 μg/ml. Selected clones were validated by PCR and whole genome sequencing to rule out ectopic mutations using Illumina paired-end libraries as described previously. Sequences were deposited in the European Nucleotide Archive with accession number ERR7179810.

For genetic complementation, the *motAB* locus (locus tags ODI_R2121 and ODI_R2122, including the promoter region) was amplified by PCR using primers motAB-Fwd-KpnI and motAB-rev-SacI (Table S2) and ligated into plasmid pBBR1MCS-3 after restriction with enzymes SacI and KpnI (NEB). Ligation products were transformed into *E. coli* Top10 by electroporation. Constructs were verified using PCR and Sanger sequencing. Plasmids were introduced into *O. dioscoreae* by electroporation. Briefly, 1 mL overnight cultures of *O. dioscoreae* were washed 3 times in sterile ultrapure water and resuspended in 40 μL. About 0.5 μg of plasmid DNA were mixed with the cell suspension and transferred to ice-cold 1mm gap cuvettes (Bio-Rad). Cells were electroporated in a Bio-Rad Gene Pulser Xcell system using settings: 1.8 kV voltage, 25μF, 200 Ω. Transformants were selected on TSA medium supplemented with tetracycline (20 μg/L).

### Transmission electron microscopy

Samples were fixed in 2% (v/v) glutaraldehyde (EMS) + 0.5% (v/v) paraformaldehyde (EMS) in a 50 mM sodium buffer, pH 7.2 at room temperature and under vacuum. After four hours, the fixative solution was refreshed and samples were kept at 4°C for 26 days. Samples were rinsed twice in 50 mM cacodylate sodium buffer (pH 7.2) and postfixed in 2% (v/v) osmium tetroxide in water for 1.5 hours at room temperature in darkness. Samples were rinsed three times in water and dehydrated using a graded ethanol series (10%-100%, 10% increments). Samples were then incubated in propylene oxide (PO) (EMS)) for 2 times 1 hour and infiltrated in Epon using a PO/Epon series over multiple days at 4°C. Samples were embedded in flat embedding molds and polymerized for 48 hours at 60°C. Thin sections of 1 μm were cut using a Leica Ultracut E Reichert and contrasted using Uranyless and lead citrate (Delta Microscopies, France). Samples were viewed using a Hitachi HT7700 electron microscope.

### Scanning electron microscopy

Samples were fixed in 2.5% (v/v) glutaraldehyde in 50 mM cacodylate sodium buffer (pH 7.2) for 3 hours at RT and transferred to 4°C for 2 days. Samples were dehydrated using a graded ethanol series. The samples were dried using a critical point drier (Leica EM CPD 300) using CO_2_ as transitional medium. A platinum coating was applied and samples were examined using a FEG FEI Quanta 250 electron microscope.

### Light Microscopy

Samples were fixed in 4 % (v/v) formaldehyde in PEM buffer (100 mM 1,4-piperazinediethanesulfonic acid, 10 mM MgSO_4_, and 10 mM ethylene glycol tetra-acetic acid, pH 6.9) and rinsed in water. Samples were washed in PBS (Na_2_HPO_4_ 0.148 g, KH_2_PO_4_ 0.043 g, NaCl 0.72 g, NaN_3_ 0.9 g in 100 mL distilled water, pH 7.1) and dehydrated using a graded ethanol series (30, 50, 70, 85, 100 % (v/v)). Samples were polymerised in LR White acrylic resin (medium grade, London Resin Company, UK) using polypropylene capsules at 37 °C for three days. Semi-thin sections of 350 nm were cut using Leica UC6 ultramicrotome (Leica Microsystems, Vienna) equipped with a diamond knife. Sections were collected on polylysine-adhesion slides (Carl Roth, Germany). Sections were stained with 1% (w/v) toluidine blue O (Merck, Germany) in 1% Na_2_B_4_O_7_ for 20 seconds at 50°C, rinsed with dH_2_O and mounted in DePeX.

Samples stained with Calcofluor and Auramine O were processed as followed: Wild-type acumens were fixed in 4% paraformaldehyde in PBS at 4°C overnight, washed twice in PBS and cleared by subsequently incubating samples in clearing solutions for one week at 37°C. The first solution contained 5% v/v glycerol + 10% v/v sodium deoxycholate + 10% v/v urea + 10% v/v xylitol and urea and xylitol concentrations increased to 20% and 30% in week 2 and week 3, respectively. Cleared samples were stained overnight at 4°C in 0.01% Calcofluor and 0.01% Auramine O.

For vibratome sectioning, samples were enclosed in 8 % agarose, glued upon the cutting stage using superglue (Roticoll 1, Carl Roth, Karlsruhe, Germany) and cut into 30 μm thick sections with a vibrating microtome (HM650V, Thermo Fisher Scientific, Waltham, MA, USA). Sections were stained with 0.5% (w/v) astra blue, 0.5% (w/v) chrysoidine and 0.5% (w/v) acridine red for 3 minutes, rinsed in water, dehydrated with isopropyl alcohol and mounted in Euparal (Carl Roth, Karlsruhe, Germany). All sections were observed using a Nikon Eclipse Ni-U bright field microscope equipped with a Nikon DS-Fi1c camera.

To visualize mCherry tagged *O. dioscoreae* (R71417) in the shoot tips, fresh plant samples were sectioned with a razor blade and imaged using a laser scanning confocal microscope (Leica TCS SP2). LAS X software was used to process the images.

### Estimation of infection bottleneck

Bacterial strains (R-67170 and R-71416) were cultured in TSB medium. Bacterial cultures in the exponential phase of growth were centrifuged (5000 rpm, 10 min) and washed twice with 0.4% NaCl. Cell suspensions were normalized to OD_600nm_ = 0.2. Suspensions of R-71416 (GFP-tagged and resistant to gentamycin) were serially diluted with suspensions of the non-tagged strain to yield different concentrations of target strain (1:1, 1:10, 1:100, 1:1000, 1:10 000, 1:100 000) at a constant OD. These suspensions were used to inoculate aposymbiotic plants as described above. Per condition, 5 plants were inoculated. Plants were left at 28°C, 16h of light. After 5 weeks, acumens of young leaves were ground in 100μl sterile 0.4% NaCl as described above and serial dilutions were plated out on selective (TSA medium supplemented with nalidixic acid 30 μg/ml and gentamycin 50 μg/ml) and non-selective (TSA medium supplemented with nalidixic acid 30 μg/ml) medium as described above.

### In vitro motility test

Bacteria of interest were grown in liquid culture in TSB medium and 5 μl of overnight cultures were spotted on motility agar medium: pancreatic digest of casein Bacto peptone (10g/L), meat extract (3g/L), sodium chloride (5g/L) and agar: (4g/L), triphenyltetrazolium chloride (TTC) 0.05g/L. Plates were incubated at 28°C and the bacterial halo was measured after 48 hours.

### Measurement of gene expression

Apical buds and leaf acumens were ground in liquid nitrogen. RNA samples (four biological replicates per sample) were isolated using the RNeasy Plant Mini Kit (Invitrogen) with DNAse treatment following the manufacturer’s recommendations. Ribonucleic acid was quantified using a NanoDrop Spectrophotometer ND-100 (NanoDrop Technologies, Wilmington, DE, USA) and integrity was evaluated with a Bioanalyzer 2100 (Agilent Technologies, Santa Clara, CA, USA). Reverse transcription was performed with 2 μg of total RNA using the Reverse transcriptase Superscript II (Invitrogen) and random hexamer primers (Eurofins Genomics, Germany) for bacterial transcript quantification. Quantitative PCRs were conducted with SybrGreen (Roche) on 384-well plates using a LightCycler 480 (Roche) following manufacturer recommendations and the primers shown in Table S2. The *gyrB* encoding gene was used as an internal standard for sample comparisons. The specificity and efficiency of the amplification were verified by analyses of melting curves and standard curves, respectively. The 2^-ΔΔCt^ method was used for the calculation of relative expression (35).

## Results

### Anatomy of the *D. sansibarensis* leaf gland and relationship with the symbiotic bacteria

To investigate the distribution of the symbiotic bacteria in *D. sansibarensis*, we dissected various surface-sterilized organs and tissues and counted colonies of *O. dioscoreae* after maceration and serial dilution plated on TSA medium (Table 1). The acumens on *D. sansibarensis* leaves contain the highest number of viable bacteria, with 2.31 x 10^11^ cfu/g on average (Figure 1, Table 1). Cross-sections of the forerunner tip showed from 2 kidney-shaped glands, and up to 6 glands per acumen in large leaves (Figure 1B). Glands run along the entire length of the acumen and are lined by a cuticle (Miller & Reporter, 1987). Glands are closed at the adaxial side, where a remaining suture is apparent (Figure 1B). A thick outer layer, which stains intensely with Auramine O, lines the inside of the glands. Auramine O is a lipophilic fluorescent dye with affinity for regions containing acidic and unsaturated cuticle waxes (36). This cuticle layer forms a physical barrier between the mesophyll and the lumen of the gland (Figure 1C). Long vermiform trichomes project into the lumen of the gland (Figure 1C), which also contains a high density of bacteria (Figure 1B, Figure 2A). Trichome cells contain multiple vacuoles and vesicles, indicating intense cytotic activity (Figure 2A). In addition, bacteria display a thick, electron-lucent capsule with visible membranous projections (Figure 2B). Trichome cells close to the bacteria-filled lumen contain Golgi, endoplasmatic reticula and numerous vesicles (Figure 2C). Some vesicles are seen merging with the plasma membrane indicating cytotic activity (Figure 2D). At this interface between trichome head cells and bacteria, the electron-dense cuticle presents small gaps (Figure 2C). We did not observe structures resembling bacteria inside plant cells, suggesting a strict extracellular lifestyle. Apical and lateral buds also showed high levels of endophyte colonization. However, leaf lamina and stems contain nearly undetectable quantities of bacteria (Table 1). This suggests that, outside of the leaf gland, bacteria only associate with organogenic tissues. We also detected bacteria inside bulbils, the main form of propagule of *D. sansibarensis*. Although difficult to detect by fluorescence microscopy, bacteria may be present in the intercellular spaces of the bulbil growth center, from which new shoots emerge after germination (Table 1, Figure S2).

**Figure 1:**
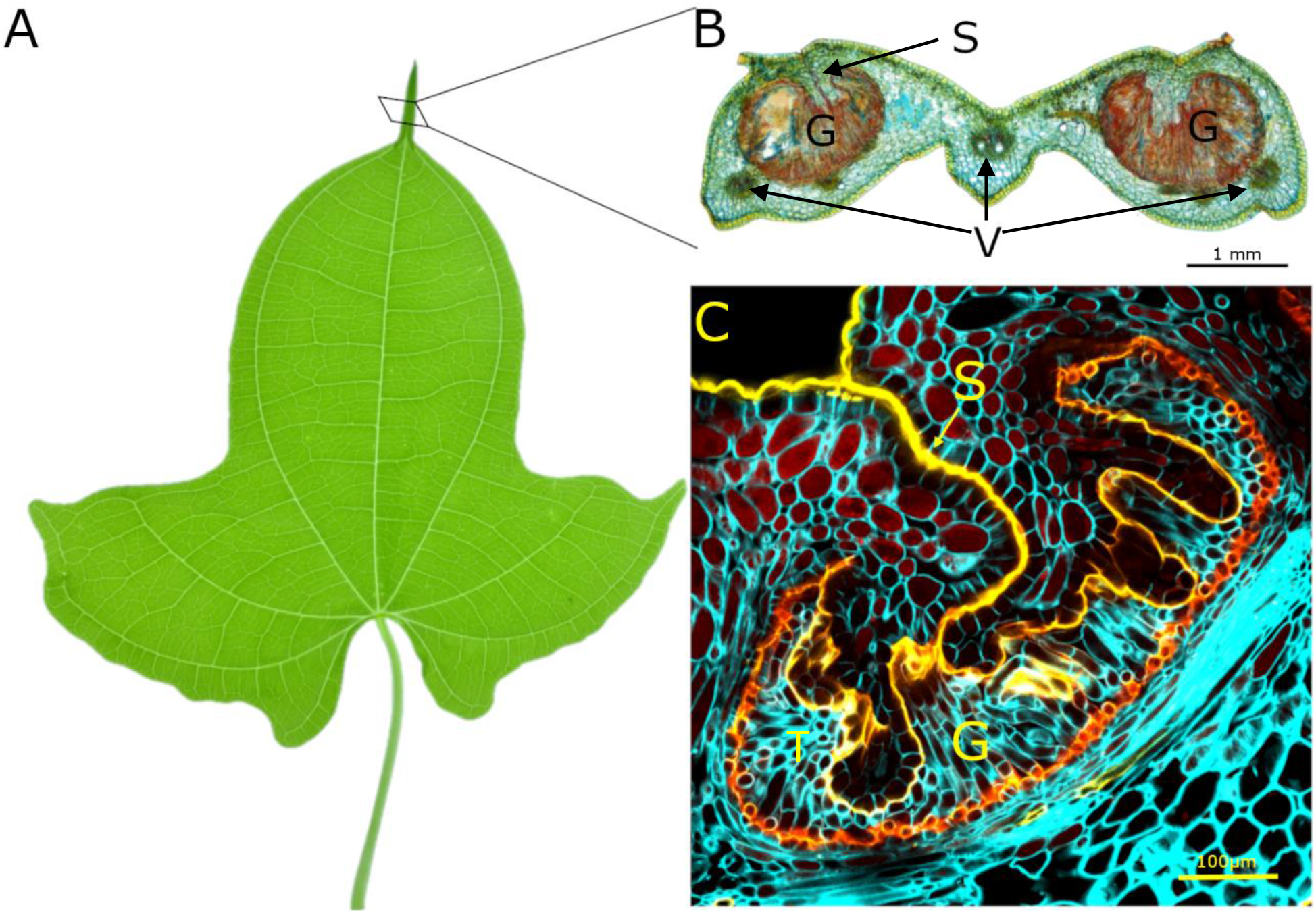
Anatomy of the *Dioscorea sansibarensis* leaf and acumen. A: Juvenile leaf from *D. sansibarensis*. Juvenile leaves are lobed and evolve to heart-shaped leaves in adult plants. Adult leaves can measure up to 46 centimeters long by 58 wide with an acumen at the distal side measuring up to 6 cm. B: An acumen cross-section. Leaves in all developmental stages contain *O. dioscoreae* in the acumen. In the acumen, two glands (G) that are filled with trichomes (T) and bacteria (stained by acridine orange) residing in mucus can be distinguished. The glands are closed at the adaxial side with a seam (S) running along the long axis of the acumen. Around the glands, several vascular bundles can be found (V). C: Auramine staining of the acumen shows the thick cuticle (yellow) surrounding the gland (G) that closes up at the adaxial side into a seam (S) and forms a physical barrier to the symbiont (not visible). Plant cell walls are stained with calcofluor white.

**Figure 2.**
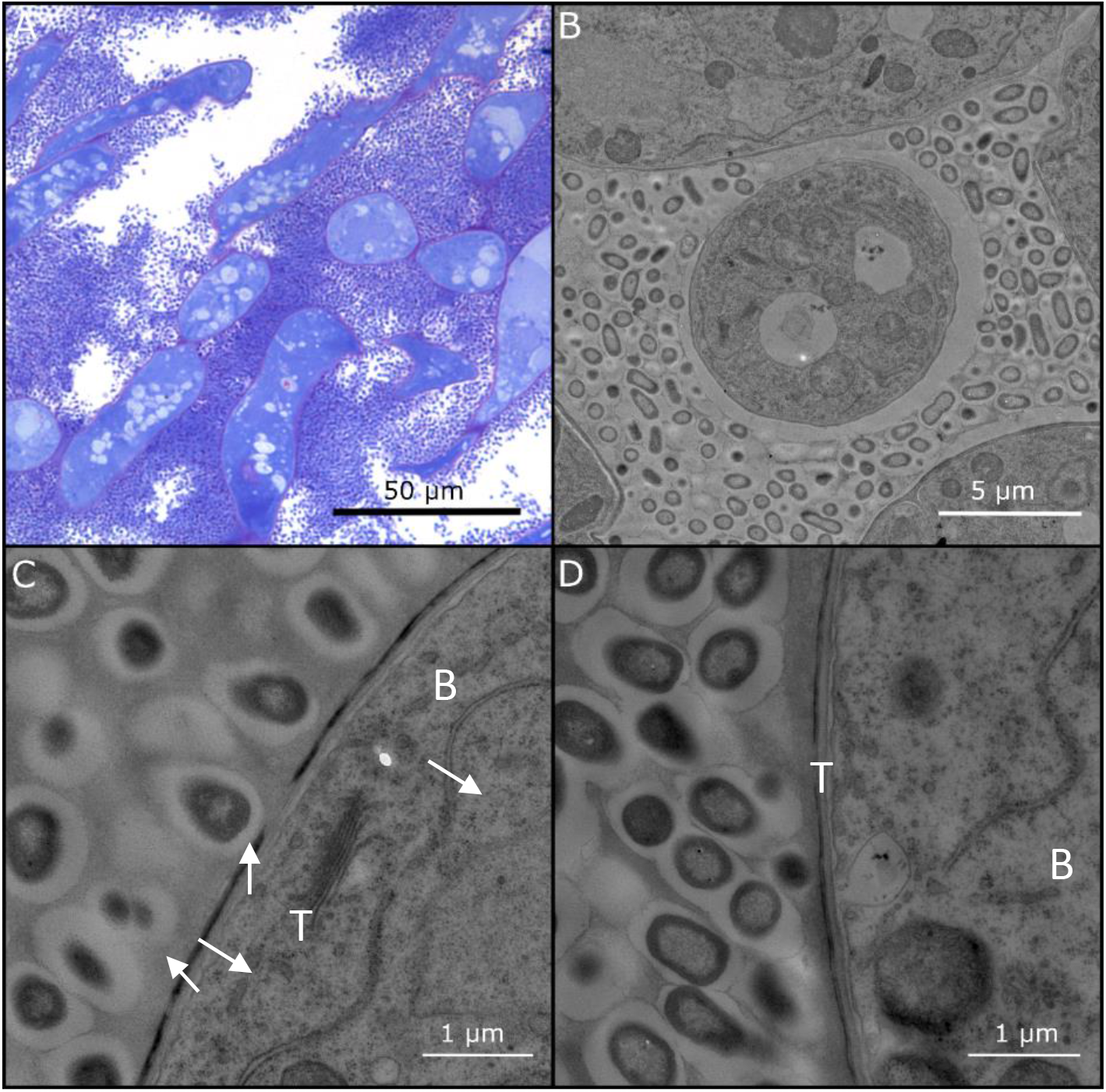
Structure of the trichome-bacteria interface in the symbiotic leaf gland. A. Light microscopic image of a cross section of the fore-runner tip of *D. sansibarensis*. Trichome cells (T) are densely stained, and multiple vesicles are visible (arrows). Around trichomes, many bacteria (B) reside in a mucilage layer. B. TEM photograph showing the cavity in a symbiotic gland. Trichomes (T) are surrounded by encapsulated bacteria (B) that fill the lumen. C. TEM image of the interface between the bacteria (B) and a trichome (T) shows the presence of multiple vesicles (white arrows), endoplasmatic reticulum (ER) and Golgi (G). Multiple gaps in the electron dense layer are apparent (hollow arrows). D. Close-up of the trichome cuticle shows a vesicle (V) merging with the plasmolemma, suggesting cytotic activity.

**Table 1.** Bacterial symbiont colonization in different tissues. Dissected and homogenized *D. sansibarensis* meristematic tissues were plated out and quantified. Of each tissue, 5 samples were taken and plated out, 95% confiden ce intervals are given.

| Tissue | CFU | Weight (mg) |
| --- | --- | --- |
| Leaf gland | $3.20 \times 10^{10} \pm 3.94 \times 10^7$ | 138.4 |
| Apical bud | $6.27 \times 10^4 \pm 4.23 \times 10^2$ | 9.3 |
| 1 cm<sup>2</sup> leaf surface | 0 | 26.1 |
| Bulbil growth centre | $7.83 \times 10^4 \pm 6.61 \times 10^2$ | 43.2 |
| Lateral bud | $3.53 \times 10^4 \pm 1.53 \times 10^2$ | 24.8 |
| Stem under apical bud | $4.87 \times 10^1 \pm 7.2 \times 10^1$ | 7.65 |

### Development of the symbiotic gland

These observations indicate that glands are an important site of exchange between the symbiotic partners. To understand how the symbiotic bacteria colonize the newly formed leaf glands, we studied the development of the gland in the apical bud (Figure 3). The leaf acumen, sometimes called a forerunner tip (37), is the first leaf structure formed as leaf primordia emerge. At later stages, the tip of the leaf folds, with margins meeting in the center to form a chamber (Figure 3B-C). Each apical bud contains 5-6 primordial leaves, of which only the three oldest develop a primordial forerunner tip (Figure 3D-F). The adaxial side of leaves displays high densities of glandular trichomes (Figure S3). The abaxial side also presents glandular trichomes, albeit in fewer numbers (Figure S3). Few visible bacteria are embedded in mucus associated with glandular trichomes at the adaxial side of young developing leaves (Figure S4), in an enclosed space delineated by the youngest leaf pair and the apical meristem which is reminiscent of the leaf enclosed chamber of *Psychotria punctata* (23)(Figure 3A). We did not observe the same shape of glandular trichomes in the closed acumens. Instead, vermiform glandular trichomes fill the gland together with mucus and the bacterial symbiont. Together, this indicates that bacteria originate from diffuse colonies near shoot meristems and colonize the symbiotic acumens as soon as the structure emerges.

**Figure 3.**
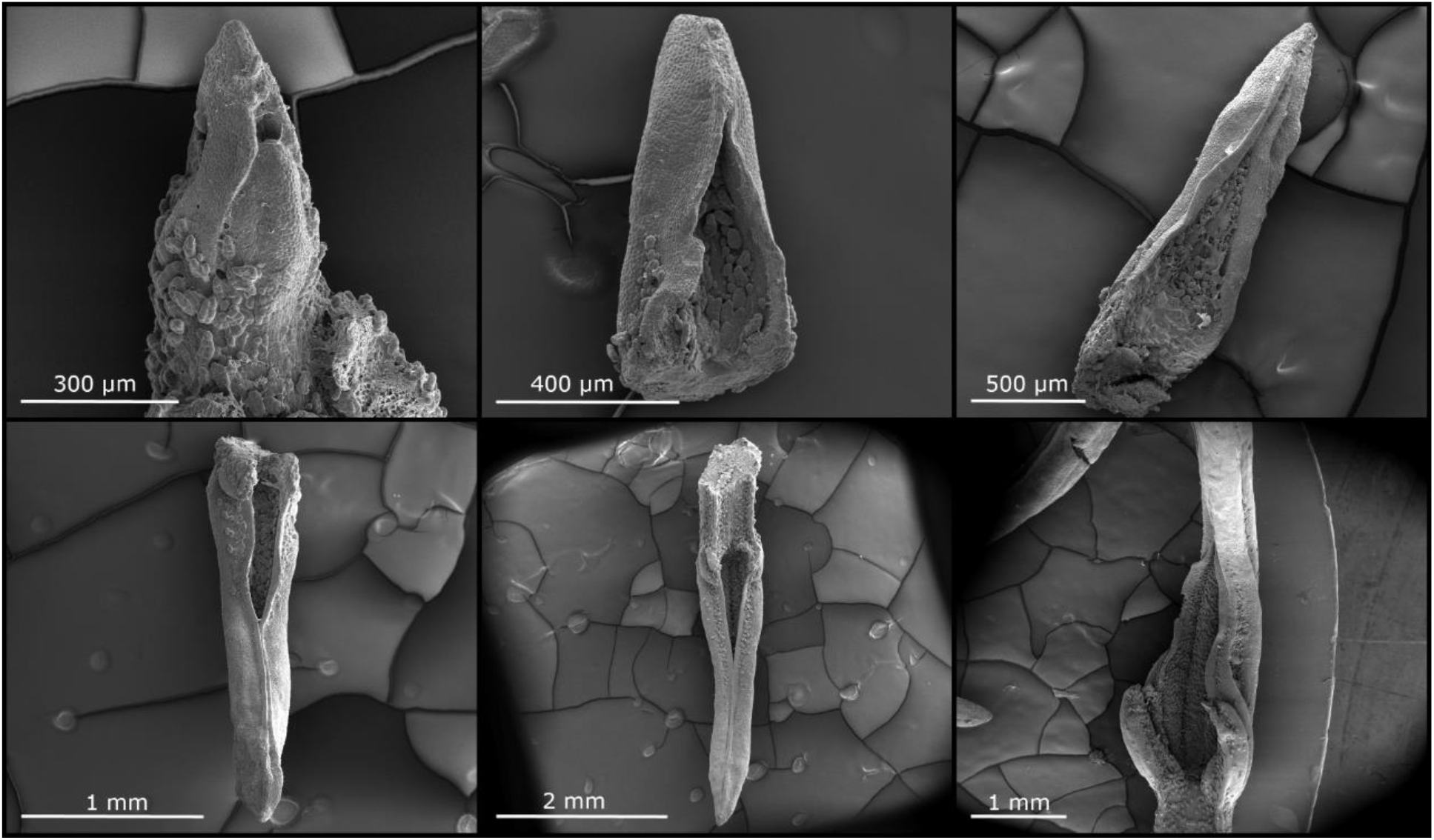
Early development of the *D. sansibarensis* leaf and fore-runner tip in the apical bud. A. Apical meristem surrounded by a leaf primordium. B. Second youngest leaf of the shoot tip, with many trichomes visible at the adaxial side and an open distal tip. C. Third youngest leaf of the shoot tip, where the acumen is starting to form, yet remains open. An abundance of trichomes and mucus can be seen at the adaxial surface. D-E-F: the third to last, second to last and last leaf of the shoot tip, respectively. The acumen progressively closes along the long axis and the leaf lamina starts to unfold slowly at the proximal side. Trichomes and mucus are abundant throughout development.

### The shoot tip is a symbiotic hub

We developed a symbiont-replacement assay to visualize the journey of *O. dioscoreae* from the apical bud to the leaf gland. To inoculate the plant with exogenous *O. dioscoreae*, we first designed a method to generate aposymbiotic plants to eliminate spatial competition. Initial attempts to obtain aposymbiotic plants by growing surface-sterilized bulbils in sterile medium as in Miller and Reporter (Miller & Reporter, 1987) consistently resulted in plants colonized by wild-type *O. dioscoreae* (data not shown). Instead, we obtained aposymbiotic plants by submerging node cuttings in a mixture of antibiotics in plant growth medium. Aposymbiotic plantlets were then inoculated by depositing cell suspensions of *O. dioscoreae* strains expressing GFP or mCherry (strains R-71416 and R-71417, respectively) onto the apical bud in otherwise sterile conditions (Figure S1). Glands of new leaves, which emerged above the point of inoculation, exclusively contained tagged bacteria, while older leaves below did not. This indicates that bacteria colonize symbiotic tissue during early development near the shoot meristem, but do not spread to older tissue via apoplastic or symplastic routes. At the apical bud, bacteria seem to adhere to the trichomes on primordial leaves (Figure 4).

**Figure 4.**
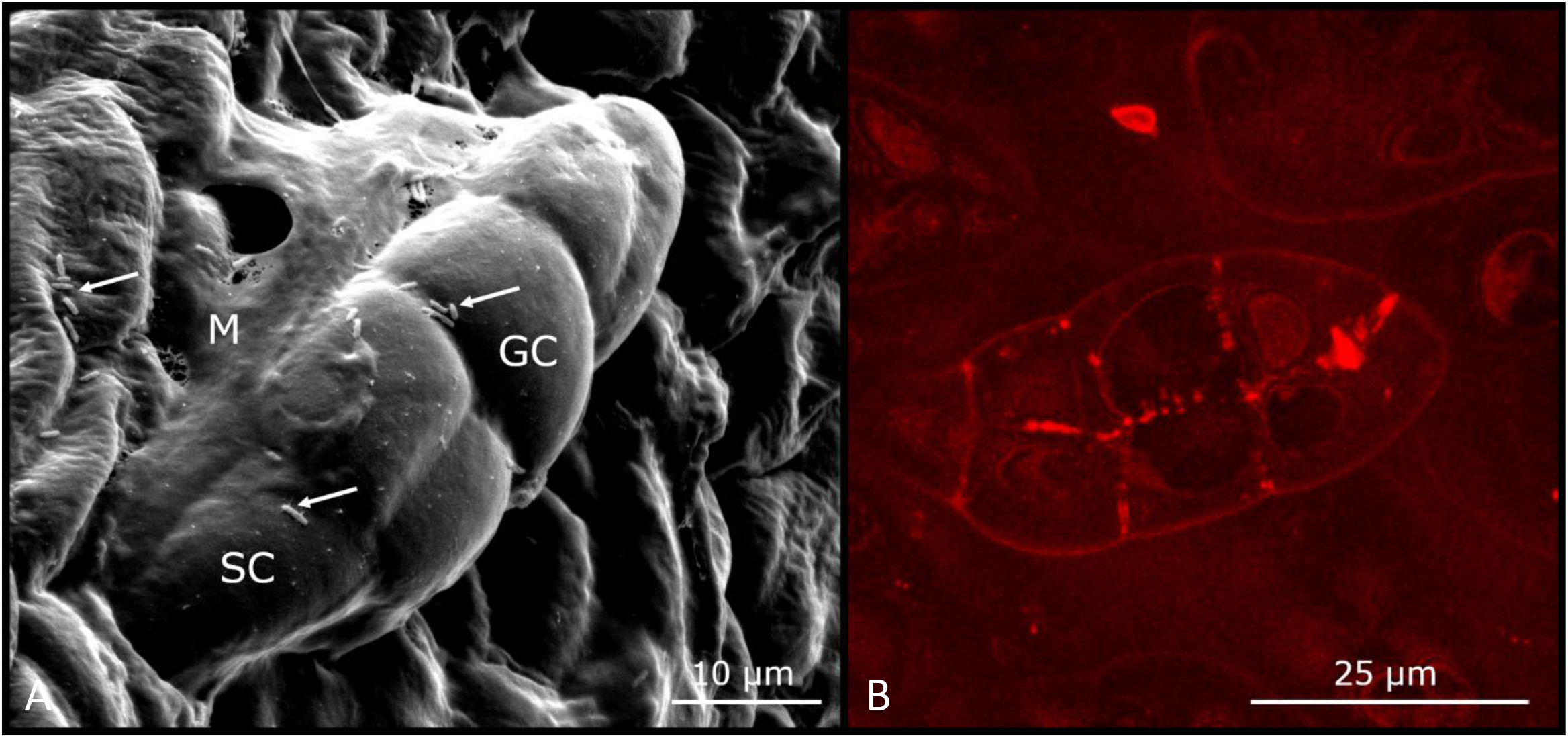
*O. dioscoreae’s* habitat in the shoot tip. Scanning electron (left) and confocal (right) microscopy pictures of leaf primordia in the shoot tip show *O. dioscoreae* colonizing the glandular trichomes. A. Trichomes in the apical bud consist of one stalk cell (SC) and 5 or 6 glandular cells (GC). Bacteria (arrow) and mucus (M) surround the trichomes. B. Confocal microscopy of glandular trichomes in the shoot tip, showing association with mCherry-tagged *O. dioscoreae*.

To investigate if bacteria are transferred to the next generation, we inoculated aposymbiotic plants with mCherry-tagged *O. dioscoreae* R-71417. After 5 weeks of growth in gnotobiotic conditions, we transferred the plants to open pots filled with soil. We harvested bulbils of the plants that survived the transfer to open pots at the end of the growing season and planted the bulbils in soil. Plants germinated from bulbils all contained fluorescent *O. dioscoreae* in their forerunner tips, demonstrating that presence of *O. dioscoreae* at the apical bud is sufficient for the colonization of plant tissues, including reproductive structures such as bulbils. These results show that the bacterial symbiont is transmitted through the bulbils. Unfortunately, we were not able to establish if sexual reproductive structures also contain symbiotic bacteria because *D. sansibarensis* rarely flowers in the wild, and never in cultivation (38).

Although artificial, our symbiont-replacement assay also shows that horizontal acquisition of *O. dioscoreae* is possible. To gain a better understanding of how likely exogenous bacteria are to enter the apical bud, we infected aposymbiotic plants with mixed cell suspensions of GFP-tagged *O. dioscoreae* R-71416 and a wild type *O. dioscoreae* R-71412 in ratios ranging from 1:1 to 1:10^5^, for a total number of approximately 2 x 10^5^ cells per inoculum. Leaf glands were harvested from plants grown in gnotobiotic conditions at 5 weeks post-infection, macerated and the contents plated on selective medium and non-selective medium to count colonies of tagged and total bacteria, respectively. We detected GFP-tagged bacteria in only 20% of plants inoculated with a dilution factor of 1:100, and none with dilution factors above 1:1000 (Figure S5). This suggests that the number of bacteria establishing in the plant is in the low hundreds. Together with the fact that all our attempts to force ingress of exogenous bacteria in already symbiotic plants failed (data not shown), we conclude that horizontal symbiont transmission is probably a rare event, in accordance with our previous phylogenetic analyses (30).

### Populations of *O. dioscoreae* in the leaf enclosed chambers and leaf glands are physiologically distinct

As bulbils and tubers grow from modified shoot buds, we propose that the small colony of *O. dioscoreae* in leaf-enclosed chambers provides the initial inoculum for the developing leaf glands, as well as lateral meristems and the reproductive organs. We hypothesized that bacteria occurring in leaf glands may dedicate their metabolism to symbiotic functions, whereas bacteria in buds may allocate resources for multiplication and transmission. We have previously identified a set of genes in three putative operons that were highly upregulated in the leaf gland compared to axenic cultures (29). The *smp1, smp2* and *opk* genes are related to non-ribosomal peptide and polyketide synthesis, respectively, and were upregulated >150-fold in the leaf gland vs. culture (29). To test if bacterial populations at the leaf glands and at the apical buds have distinct metabolic characteristics, we measured the expression of select *smp* and *opk* genes by RT-qPCR. Expression levels of *smp* and *opk* transcripts were at least 10-fold lower in apical bud bacteria compared to leaf gland (Table 2). This is likely an underestimation, since transcript levels of target genes were below detection levels in some apical bud samples (Table S3).

**Table 2.**
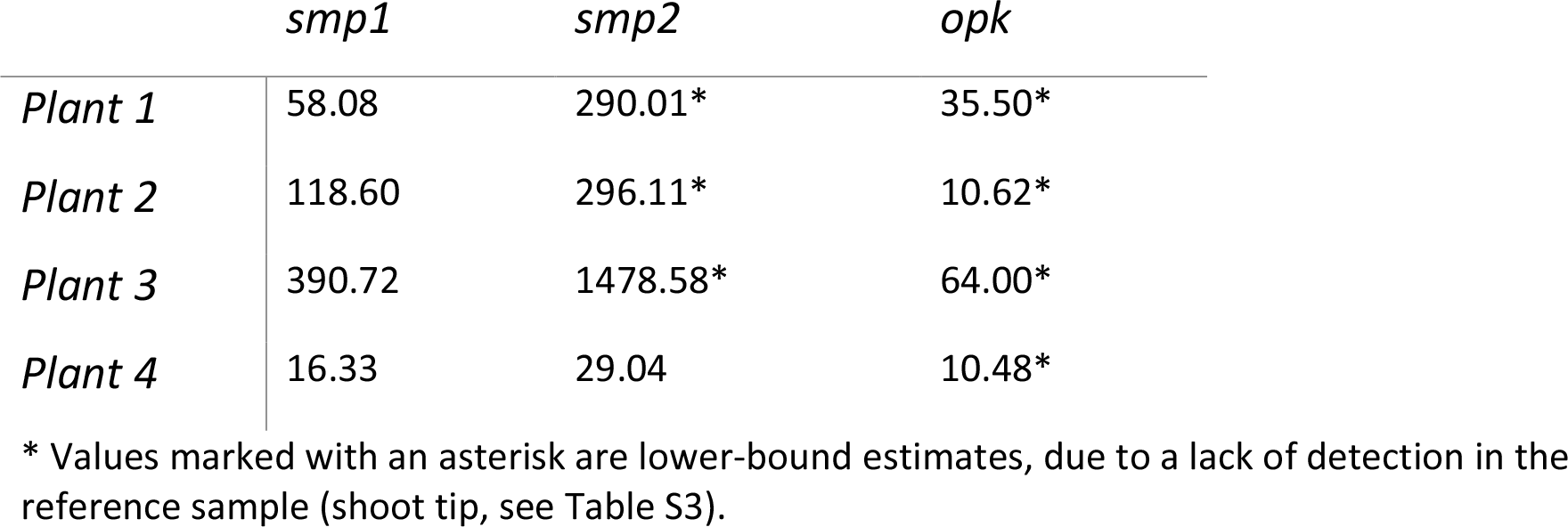
Differential regulation of *O. dioscoreae* putative secondary metabolism in leaf gland vs shoot tip by quantitative RT-PCR. Acumen and shoot tip RNA samples were collected during the day. Primers nrp 88-89, pqqc 90-91 and KASII 82-83 correspond to a representative gene from the *smp1* (ODI_R1490), *smp2* (ODI_R1505) and *opk* (ODI_R2249) gene clusters, respectively. Fold changes (2^-ΔΔCt^) in transcript abundance are given in the leaf gland *vs*. the apical shoot tip.

### Motility is dispensable for host colonization, but necessary for horizontal transmission

Motility is often required by plant pathogens and symbionts to colonize their hosts (39–43). Our observation that *O. dioscoreae* do not colonize leaf glands below the inoculation point however suggests that movement of bacteria within the plant is limited. Moreover, obligate *Burkholderia* leaf symbionts of Rubiaceae and Primulaceae lack flagella, suggesting that motility is not essential for within-host spread or vertical transmission in leaf symbiosis (16). To test whether flagellar motility is required for colonization of *D. sansibarensis*, we generated strain *O. dioscoreae* TA01 by allelic exchange with a copy of a *motB* homolog (locus tag ODI_R2122) interrupted by a kanamycin resistance cassette. MotB is a component of the flagellar motor complex and is essential for motility (44). We confirmed that MotB is involved in motility in *O. dioscoreae* by measuring the halo of colonies spotted onto soft motility agar. The colony diameter of strain R-71417 (WT) on motility agar was 6.03 ± 1.11 cm (95% confidence interval) while strain TA01 (*ΔmotB*) was unable to move beyond the initial spot on the agar (colony diameter of 0.93 ± 0.05 cm (95% C.I.). The complemented strain TA01 *motB^+^*, showed intermediate levels of motility with a colony diameter of 2.17 ± 1.23 cm (95% C.I.). Importantly, we did not notice a difference in growth rates between strains TA01 and R-71417 (data not shown). To test the effect of impaired motility *in planta*, we introduced strains R-71417 or TA01 into aposymbiotic plants. After inoculation and incubation for five weeks, leaf glands were macerated, and the contents plated out on selective media to allow for selective counting of strain R-71417 or TA01. Colonization rates were high across all conditions, with bacteria in the leaf glands of 59 out of 68 plants (one to four leaves checked per plant). The success rate of inoculations with R-71417, TA01 or TA01 *motB+* did not differ significantly in single inoculations, with 75%, 66.67%, and 80% of plants successfully colonized, respectively. Furthermore, bacterial densities inside leaf glands did not differ significantly between plants inoculated with parental strain R-71417, TA01, or TA01 *motB^+^* (Student T-test *p*-value > 0.05, Figure 5A). Altogether, this indicates that flagellar motility is not required for host colonization. However, the inoculum used in our assays contained a large excess of bacteria and these results may not reveal subtle differences in colonization fitness between the strains. To test whether non-motile strains are outcompeted by motile strains in our assay, we performed co-inoculations of aposymbiotic plants with strain TA01 and R-71417 in 1:1 ratio. Bacterial densities of strain TA01 inside leaf glands were between 10 to 10^10^ times lower than R-71417, with a median competitive index of 1.0 x 10^-5^ (Figure 5B). Moreover, we performed a complementation experiment by co-inoculating strains TA01 *motB^+^* and R-71417 pBBR1MCS. The expression of a functional copy of *motB in trans* significantly raised the competitive index of strain TA01 *motB^+^*, bringing it to a median value of 1.0 x 10^-3^ (Two-sided Wilcoxon rank sum test *p* < 0.05). These data thus show that flagellar motility is not required for plant colonization, but may facilitate horizontal transmission and host-switching.

**Figure 5.**
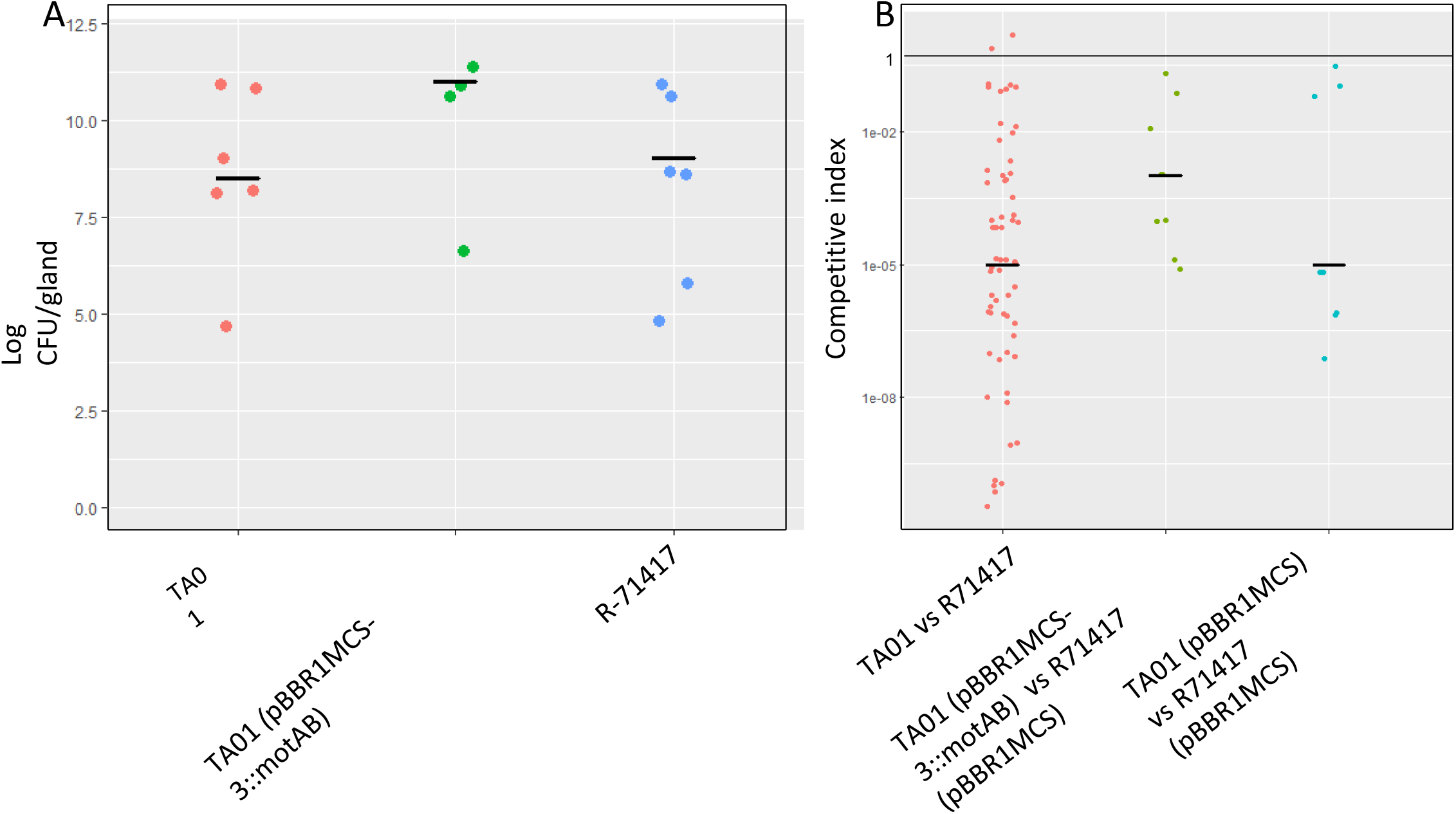
Quantification and competitive index of *O. dioscoreae* strains *in planta*. A. colonization quantification of acumens after single inoculation on the apical bud, one leaf per biological sample. Newly grown acumens were macerated and plated out, growth was quantified by colony counting. Single inoculation with *O. dioscoreae*, either mCherry-tagged strain (R-71417), the motility impaired mutant (TA01) or the complemented strain (TA01 pBBR1MCS-3::*motAB*). There is no significant difference in average bacterial densities between the 3 conditions (pairwise two-sided Student’s T-test *p* > 0.05). B. Competitive index of O*. dioscoreae* in co-inoculations of motility-impaired mutant vs. parental strain. Aposymbiotic plants were co-inoculated with a 1:1 mix of motility impaired mutants (strain TA01) and the parental strain (R-71417). As a control, a 1:1 mix of a complemented motility mutant (TA01 pBBR1MCS-3::*motAB*) with the parental strain containing the empty plasmid used for complementation (R-71417 pBBR1MCS) was inoculated into aposymbiotic plants. Lastly, to control for the effect of the empty plasmid, strain TA01 pBBR1MCS was co-inoculated with strain R-71417 pBBR1MCS.

## Discussion

The unusual tractability of the *D. sansibarensis/O. dioscoreae* symbiosis makes this association a valuable model system to study the determinants of vertical transmission of plant microbiota, as well as the molecular mechanisms governing the specificity of association of plants with bacteria at the leaf surface. In this work, we show that *O. dioscoreae* symbiotic bacteria are housed in specialized structures at the tip of the leaves, formed by the folding of the leaf margins. These glands hold high densities of bacteria (up to 10^11^ CFU/g) which are separated from the epidermis by a cuticle layer. Several lines of evidence indicate that the large numbers of trichomes which project inside the leaf gland may play an essential role in the interaction with the bacterial symbionts. The cuticle layer and cell wall appear thinner in the area directly in contact with the bacteria, with zones of discontinuity in the electron-dense layer (Figure 2). The plant cuticle acts as a diffusion barrier for water and hydrophilic compounds (45), and gaps in the cuticle layer may enable the diffusion of water-soluble and ionic solutes (46). The presence of numerous vesicles in trichome head cells supports the hypothesis that trichomes act as a major interface between the symbiotic partners. These specialized trichomes are possibly involved in the delivery of nutrients to the bacterial symbiont as well as the uptake of metabolites of bacterial origin. The genome of *O. dioscoreae* does not contain genes coding for secreted polysaccharide-degrading enzymes, and lacks a functional glycolysis, Entner-Doudorroff pathway or oxidative branch of the pentose phosphate pathway. However, *O. dioscoreae* isolates display leucine arylamidase activity (28), indicating that the bacteria have the ability to mineralize organic nitrogen in peptide bonds (47). Trichome secretions may thus be at least partly responsible for the mucous substance surrounding the bacteria in the leaf gland. Although of unknown chemical composition, this mucus may play a direct, possibly dual role as biofilm matrix component and source of complex nutrients.

Although the leaf gland is the most striking feature of the symbiosis, *O. dioscoreae* inhabits other aerial tissues. The distribution of bacteria within the host is however not random. Somatic tissues like stems or leaf lamina contain very few bacteria, but shoot organogenic tissues such as apical and lateral buds, as well as vegetative propagules (bulbils), consistently contained symbiotic bacteria. Similar to within the leaf glands, bacteria are found in a mucus, which surrounds putative secretory trichomes. Although the bacterial colonies near the shoot apical meristem are more diffuse, bacteria grow within a structure analogous to the leaf-enclosed chamber, previously described in leaf nodulating *Psychotria* species (20, 23). This leaf-enclosed chamber lacks the striking compartmentalization seen in the leaf gland. *D. sansibarensis* thus seems to tolerate bacteria in contact with shoot meristematic tissue, although bacterial densities in buds and growth centers of bulbils are several orders of magnitude lower than in the leaf gland (Table 1). This close proximity of bacteria at the shoot tip is surprising, since shoot meristems are often thought to be sterile (48). However, recent studies indicate that some species host specific bud-associated microbiota (49). Because shoot meristems are the hub of aerial organogenesis, tolerance of bacteria near shoot meristematic tissue may be a key feature of the *D. sansibarensis/O. dioscoreae* symbiosis that enables a permanent symbiotic association.

Inoculation of fluorescent-tagged *O. dioscoreae* at the shoot tip resulted in plants that contained bacteria in all leaves formed above the point of inoculation. Strikingly, we never found evidence of bacteria in the glands of leaves which had emerged prior to the time of inoculation. Microscopic observation also indicates that *O. dioscoreae* attaches to the trichomes of new leaves as they emerge from primordia, before the folding of the tip takes place (Figure 4). Together, this supports the view that growth and distribution of the symbiont is concomitant with leaf development, and supported by elongation after gland formation. The fact that we were able to inoculate aposymbiotic plants artificially also suggests that access to the leaf-enclosed chamber remains open after germination. However, we show that out of 2 x 10^5^ *O. dioscoreae* cells, only a few hundred successfully establish in the host after inoculation. This is indicative of stringent barriers to inoculation, similar to some symbiotic systems with horizontal transmission, for example that between the bean bug *Riptortus pedestris* and *Burkholderia* (50, 51), or between *Vibrio* and the bobtail squid (52). Whether the potential infection barriers in *D. sansibarensis* are selective to *O. dioscoreae* or if they allow ingress of other bacteria remains to be tested.

Similar to symbiotic systems with horizontal transmission, flagellar motility contributes to the colonization of new hosts (53–55). The infectivity of *O. dioscoreae motB* mutants was several orders of magnitude lower than that of reference strains (Figure 5). This suggests that flagellar motility facilitates crossing of host barriers to reach the leaf-enclosed chamber and propagate within the host. Despite this competitive disadvantage, *O. dioscoreae motB* mutants were still capable of infecting aposymbiotic plants and grew to normal densities *in planta*. Moreover, the genome of *O. dioscoreae* lacks genes for alternative types of motility, such as twitching motility (28). Interestingly, genes linked to chemotaxis or motility functions are entirely lacking from the genomes of some *Burkholderia* leaf nodule symbionts of Rubiaceae and Primulaceae (16, 56, 57). These data strongly suggest that bacterial motility is not required for within-host colonization and trans-generational transmission, but may instead facilitate horizontal transmission and host switching. Indeed, phylogenetic analysis indicate a strict vertical mode of transmission for leaf symbionts entirely lacking a flagellar apparatus (17, 30, 56). Evidence that motility appears dispensable *in planta* indicates that spread of the symbiotic bacteria from the leaf-enclosed chamber to the leaf glands perhaps relies on attachment to specific host structures within the plant. Similar modes of growth and transmission has been hypothesized for vertically-transmitted fungal endophytes of grasses: *Epichloë* hyphae attach to host cells at the shoot apical meristem and elongate simultaneously with leaf tissue, allowing asymptomatic colonization of leaves (58). The number of hyphae remains constant in tissue as leaves mature, and may be an adaptation to avoid uncontrolled proliferation and triggering of plant defenses (59). In *D. sansibarensis*, the number of bacteria remains constant in apical and lateral buds, as well as bulbil growth centers with approximately 1-7 x 10^6^ cfu/g of tissue (Table 1). Reciprocal signaling events between host and symbiont presumably control attachment and growth of the bacteria in leaf tissue, a key feature of this leaf symbiosis.

In addition to controlling bacterial proliferation, specific signals may also control expression of bacterial symbiotic functions in target tissue (60, 61). The *smp* and *opk* genes of *O. dioscoreae* encode putative enzymes of the secondary metabolism, which we hypothesized to play a central role in the leaf symbiosis (29, 30). Genes of the *smp* and *opk* putative biosynthetic gene clusters are highly expressed in the leaf gland, representing nearly 30% of all mRNA (29). However, our data reveal that bacteria in the apical buds express key *smp* and *opk* genes in much lower levels than in the leaf gland (Table 2). This difference in expression may reflect a strategy by the bacteria to maximize use of limited resources in the apical buds towards growth. We propose a model whereby two distinct populations of *O. dioscoreae* are maintained in the plant (Figure 6): bacteria in organogenic structures (e.g. apical or lateral buds) maintain synchronous growth with host tissue to serve as a mother colony. Bacteria of this “reproductive” pool have two distinct fates: serve as an inoculum for the leaf gland, and transmit bacteria to the next generation via propagules. Bacteria in the leaf gland provide the main symbiotic services to the plant via secretion of metabolites, but are at a reproductive dead-end. *D. sansibarensis* is an annual plant, with leaves senescing at the end of the season and bacteria presumably dying or at least excluded from the reproductive pool in the plant. We postulate that this division of labor between reproductive and productive symbionts may have important consequences for the evolution of leaf symbiosis, and would ensure that “cheater” bacteria, which do not provide symbiotic services to the host, do not outcompete mutualistic bacteria for access to plant reproductive structures.

**Figure 6.**
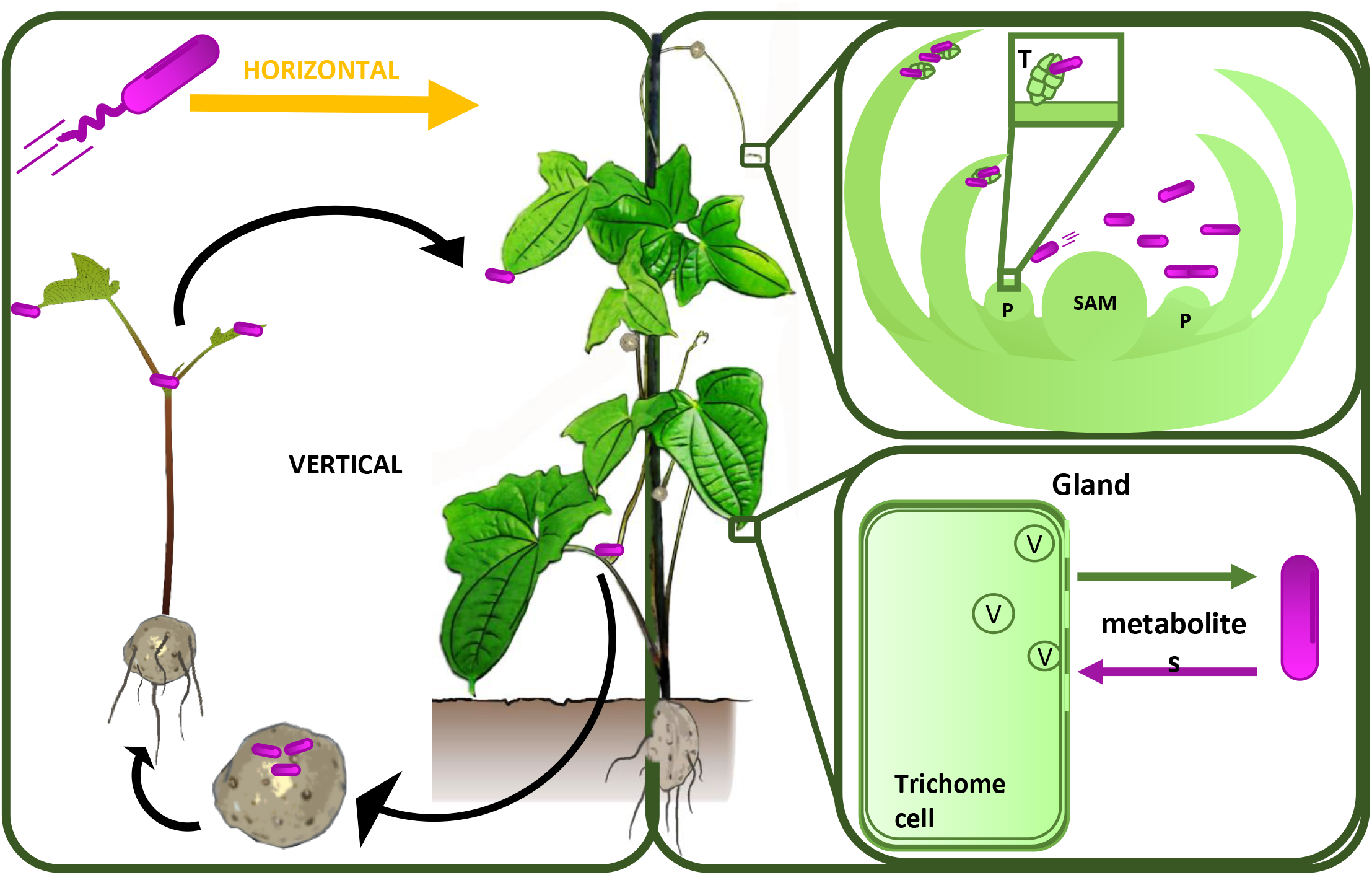
Schematic of *O. dioscoreae* transmission and functional predictions. *D. sansibarensis* harbors symbiotic bacteria (*O. dioscoreae*) which are contained within leaf glands, bulbils, and shoot apical or axillary buds. The apical bud acts as a reservoir for the symbiont to ensure colonization of newly formed aerial organs, such as the forerunner tip, lateral buds and propagules. Allocation of *O. dioscoreae* cells to bulbils ensures transmission to the next plant generation. Flagellar motility is hypothesized not being required for vertical transmission, but occasional horizontal transmission seem to at least partly rely on bacterial motility (Left panel). The main function of *O. dioscoreae* of this “reproductive” pool residing in the shoot tip may be to serve as an inoculum for the leaf gland and transmit bacteria to the next generation via bulbils. Accordingly, biosynthesis of bacterial secondary metabolites is down-regulated in the apical bud. *O. dioscoreae* attaches to trichomes (T) in the apical bud and grows synchronously with the plant (top right panel). In contrast, the fate of *O. dioscoreae* in the leaf gland may be to provide secondary metabolites to the plant, via exchange of metabolites through the permeable cell wall of specialized trichomes (lower right panel). Leaf gland bacteria are at a reproductive dead-end and are not transmitted to the next plant generation. P, leaf primordium; SAM, shoot apical meristem; V, vesicles; T, trichome.

In conclusion, we provide direct experimental evidence of vertical transmission of symbiotic bacteria in *Dioscorea sansibarensis*. Our work thus provides fundamental insights into the mechanisms governing host colonization and transmission of vertically-transmitted bacteria in plants. The unique tractability of the *Dioscorea/Orrella* association makes this an appealing model to understand mechanisms of non-pathogenic plant-bacteria interactions in the phyllosphere.

## ACKNOWLEDGMENTS

We would like to thank Thibault Sana and Bram Danneels for helpful discussion and for proofreading the manuscript. This work was supported by the UGent Special Research Fund under grant BOFSTA2017002001 to AC. AC also acknowledges support from the French National Research Agency under grant agreement ANR-19-TERC-0004-01 and from the French Laboratory of Excellence project “TULIP” (ANR-10-LABX-41; ANR-11-IDEX-0002-02). The funders had no role in study design, data collection and analysis, decision to publish, or preparation of the manuscript.

## AUTHOR CONTRIBUTIONS

TA and AC designed the research; TA, SM,FDM, OC, and MB carried out the experiments. TA, AC, OC, OD and PW analyzed data; TA and AC wrote the manuscript with input from all authors.

## DATA AVAILABILITY

The datasets generated and/or analyzed during the current study are available in the European Nucleotide Archive repository, with the following accession number: ERR7179810.

## References

1. Chen R, Wang Z, Chen J, Jiang L-Y, Qiao G-X. 2017. Insect-bacteria parallel evolution in multiple-co-obligate-aphid association: a case in Lachninae (Hemiptera: Aphididae). Sci Reports 2017 71 7:1–9.

2. Sapp J. 2002. Paul Buchner (1886-1978) and hereditary symbiosis in insects. Int Microbiol. Sociedad Espanola de Microbiologia.

3. Xu T-T, Chen J, Jiang L-Y, Qiao G-X. 2018. Historical and cospeciating associations between Cerataphidini aphids (Hemiptera: Aphididae: Hormaphidinae) and their primary endosymbiont Buchnera aphidicola. Zool J Linn Soc 182:604–613.

4. Moran NA, McCutcheon JP, Nakabachi A. 2008. Genomics and evolution of heritable bacterial symbionts. Annu Rev Genet. Annu Rev Genet.

5. McCutcheon JP. 2021. The Genomics and Cell Biology of Host-Beneficial Intracellular Infections. Annu Rev Cell Dev Biol 37:115–142.

6. Bennett GM, Moran NA. 2015. Heritable symbiosis: The advantages and perils of an evolutionary rabbit hole. Proc Natl Acad Sci U S A 112:10169–10176.

7. Martin FM, Uroz S, Barker DG. 2017. Ancestral alliances: Plant mutualistic symbioses with fungi and bacteria. Science (80-) 356.

8. Fisher RM, Henry LM, Cornwallis CK, Kiers ET, West SA. 2017. The evolution of host-symbiont dependence. Nat Commun 8:15973.

9. Sachs JL, Skophammer RG, Regus JU. 2011. Evolutionary transitions in bacterial symbiosis. Proc Natl Acad Sci U S A 108 Suppl:10800–7.

10. Bright M, Bulgheresi S. 2010. A complex journey: transmission of microbial symbionts. Nat Rev Microbiol 8:218–230.

11. Chomicki G, Kiers ET, Renner SS. 2020. The Evolution of Mutualistic Dependence. Annu Rev Ecol Evol Syst 51:409–432.

12. Gundel PE, Rudgers JA, Whitney KD. 2017. Vertically transmitted symbionts as mechanisms of transgenerational effects. Am J Bot 104:787–792.

13. Leigh EG. 2010. The evolution of mutualism. J Evol Biol. J Evol Biol.

14. Sachs JL, Mueller UG, Wilcox TP, Bull JJ. 2004. The Evolution of Cooperation. Q Rev Biol 79:135–160.

15. Pinto-Carbó M, Gademann K, Eberl L, Carlier A. 2018. Leaf nodule symbiosis: function and transmission of obligate bacterial endophytes. Curr Opin Plant Biol. Elsevier Ltd.

16. Pinto-Carbó M, Sieber S, Dessein S, Wicker T, Verstraete B, Gademann K, Eberl L, Carlier A. 2016. Evidence of horizontal gene transfer between obligate leaf nodule symbionts. ISME J 10:2092–2105.

17. Lemaire B, Smets E, Dessein S. 2011. Bacterial leaf symbiosis in Ardisia (Myrsinoideae, Primulaceae): molecular evidence for host specificity. Res Microbiol 162:528–534.

18. Lachenaud O. 2019. Revision du genre Psychotria (Rubiaceae) en Afrique occidentale et centrale. Jard Bot Meise.

19. Lachenaud O. 2013. Le genre Psychotria (Rubiaceae) en Afrique occidentale et centrale: taxonomie, phylogénie et biogéographie. Thesis, Univ Lbre Bruxelles, Bruxelles.

20. Miller IM. 1990. Bacterial leaf nodule symbiosis. Adv Bot Res Inc Adv Plant Pathol 17:163–234.

21. Lemaire B, Janssens S, Smets E, Dessein S. 2012. Endosymbiont transmission mode in bacterial leaf nodulation as revealed by a population genetic study of Psychotria leptophylla. Appl Environ Microbiol 78:284–287.

22. Lemaire B, Vandamme P, Merckx V, Smets E, Dessein S. 2011. Bacterial leaf symbiosis in angiosperms: Host specificity without Co-Speciation. PLoS One 6:e24430.

23. Sinnesael A, Eeckhout S, Janssens SB, Smets E, Panis B, Leroux O, Verstraete B. 2018. Detection of Burkholderia in the seeds of Psychotria punctata (Rubiaceae) – Microscopic evidence for vertical transmission in the leaf nodule symbiosis. PLoS One 13:e0209091.

24. Crüsemann M, Reher R, Schamari I, Brachmann AO, Ohbayashi T, Kuschak M, Malfacini D, Seidinger A, Pinto-Carbó M, Richarz R, Reuter T, Kehraus S, Hallab A, Attwood M, Schiöth HB, Mergaert P, Kikuchi Y, Schäberle TF, Kostenis E, Wenzel D, Müller CE, Piel J, Carlier A, Eberl L, König GM. 2018. Heterologous Expression, Biosynthetic Studies, and Ecological Function of the Selective Gq-Signaling Inhibitor FR900359. Angew Chemie Int Ed 57:836–840.

25. Sieber S, Carlier A, Neuburger M, Grabenweger G, Eberl L, Gademann K. 2015. Isolation and Total Synthesis of Kirkamide, an Aminocyclitol from an Obligate Leaf Nodule Symbiont. Angew Chemie 127:8079–8081.

26. Sinnesael A, Leroux O, Janssens SB, Smets E, Panis B, Verstraete B. 2019. Is the bacterial leaf nodule symbiosis obligate for Psychotria umbellata? The development of a Burkholderia-free host plant. PLoS One 14:e0219863.

27. Miller IM, Reporter M. 1987. Bacterial leaf symbiosis in Dioscorea sansibarensis: morphology and ultrastructure of the acuminate leaf glands. Plant, Cell Environ 10:413–424.

28. Carlier A, Cnockaert M, Fehr L, Vandamme P, Eberl L. 2017. Draft genome and description of Orrella dioscoreae gen. nov. sp. nov., a new species of Alcaligenaceae isolated from leaf acumens of Dioscorea sansibarensis. Syst Appl Microbiol 40:11–21.

29. De Meyer F, Danneels B, Acar T, Rasolomampianina R, Rajaonah MT, Jeannoda V, Carlier A. 2019. Adaptations and evolution of a heritable leaf nodule symbiosis between Dioscorea sansibarensis and Orrella dioscoreae. ISME J 13:1831–1844.

30. Danneels B, Viruel J, Mcgrath K, Janssens SB, Wales N, Wilkin P, Carlier A. 2021. Patterns of transmission and horizontal gene transfer in the Dioscorea sansibarensis leaf symbiosis revealed by whole-genome sequencing. Curr Biol 31:2666–2673.e4.

31. Alizadeh S, Mantell SH, MariaViana A. 1998. In vitro shoot culture and microtuber induction in the steroid yam Dioscorea composita Hemsl. Plant Cell Tissue Organ Cult 53:107–112.

32. Choi K-H, Schweizer HP. 2005. An improved method for rapid generation of unmarked Pseudomonas aeruginosa deletion mutants. BMC Microbiol 5:30.

33. Bao Y, Lies DP, Fu H, Roberts GP. 1991. An improved Tn7-based system for the single-copy insertion of cloned genes into chromosomes of gram-negative bacteria. Gene 109:167–168.

34. Fazli M, Harrison JJ, Gambino M, Givskov M, Tolker-Nielsen T. 2015. In-frame and unmarked gene deletions in Burkholderia cenocepacia via an allelic exchange system compatible with gateway technology. Appl Environ Microbiol 81:3623–3630.

35. Livak KJ, Schmittgen TD. 2001. Analysis of relative gene expression data using real-time quantitative PCR and the 2-ΔΔCT method. Methods 25:402–408.

36. Buda GJ, Isaacson T, Matas AJ, Paolillo DJ, Rose JKC. 2009. Three-dimensional imaging of plant cuticle architecture using confocal scanning laser microscopy. Plant J 60:378–385.

37. Weber O, Wilkin P, Rakotonasolo F. 2005. A new species of edible yam (Dioscorea L.) from western Madagascar. Kew Bull 60:283–291.

38. Wilkin P, Schols P, Chase MW, Chayamarit K, Furness CA, Huysmans S, Rakotonasolo F, Smets E, Thapyai C. 2005. A Plastid Gene Phylogeny Of the Yam Genus, <I>Dioscorea</I>: Roots, Fruits and Madagascar. Syst Bot 30:736–749.

39. Herrera CM, Koutsoudis MD, Wang X, von Bodman SB. 2008. Pantoea stewartii subsp. stewartii Exhibits Surface Motility, Which is a Critical Aspect of Stewart’s Wilt Disease Development on Maize. Mol Plant-Microbe Interact 21:1359–1370.

40. Tans-Kersten J, Huang H, Allen C. 2001. Ralstonia solanacearum needs motility for invasive virulence on tomato. J Bacteriol 183:3597–3605.

41. Haefele DM, Lindow SE. 1987. Flagellar Motility Confers Epiphytic Fitness Advantages upon Pseudomonas syringae. Appl Environ Microbiol 53:2528–2533.

42. Kolton M, Frenkel O, Elad Y, Cytryn E. 2014. Potential role of flavobacterial gliding-motility and type IX secretion system complex in root colonization and plant defense. Mol Plant-Microbe Interact 27:1005–1013.

43. Böhm M, Hurek T, Reinhold-Hurek B. 2007. Twitching motility is essential for endophytic rice colonization by the N2-fixing endophyte Azoarcus sp. strain BH72. Mol Plant-Microbe Interact 20:526–533.

44. Blair DF, Kim DY, Berg HC. 1991. Mutant MotB proteins in Escherichia coli. J Bacteriol 173:4049–4055.

45. Yeats TH, Rose JKC. 2013. The Formation and Function of Plant Cuticles. Plant Physiol 163:5–20.

46. Schonherr J. 2006. Characterization of aqueous pores in plant cuticles and permeation of ionic solutes. J Exp Bot 57:2471–2491.

47. Müller T, Müller M, Behrendt U. 2004. Leucine arylamidase activity in the phyllosphere and the litter layer of a Scots pine forest. FEMS Microbiol Ecol 47:153–159.

48. Handa T, Harada H. 1992. In vitro Culture of Higher Plants. Environ Control Biol 30:53–58.

49. Pohjanen J, Koskimäki JJ, Sutela S, Ardanov P, Suorsa M, Niemi K, Sarjala T, Häggman H, Pirttilä AM. 2014. Interaction with ectomycorrhizal fungi and endophytic methylobacterium affects nutrient uptake and growth of pine seedlings in vitro. Tree Physiol 34:993–1005.

50. Kikuchi Y, Yumoto I. 2013. Efficient Colonization of the Bean Bug Riptortus pedestris by an Environmentally Transmitted Burkholderia Symbiont. Appl Environ Microbiol 79:2088–2091.

51. Kikuchi Y, Ohbayashi T, Jang S, Mergaert P. 2020. Burkholderia insecticola triggers midgut closure in the bean bug Riptortus pedestris to prevent secondary bacterial infections of midgut crypts. ISME J 2020 147 14:1627–1638.

52. Wollenberg MS, Ruby EG. 2009. Population Structure of Vibrio fischeri within the Light Organs of Euprymna scolopes Squid from Two Oahu (Hawaii) Populations. Appl Environ Microbiol 75:193–202.

53. Brennan CA, Mandel MJ, Gyllborg MC, Thomasgard KA, Ruby EG. 2013. Genetic determinants of swimming motility in the squid light-organ symbiont Vibrio fischeri. Microbiologyopen 2:576.

54. Wheatley RM, Ford BL, Li L, Aroney STN, Knights HE, Ledermann R, East AK, Ramachandran VK, Poole PS. 2020. Lifestyle adaptations of Rhizobium from rhizosphere to symbiosis. Proc Natl Acad Sci 117:23823–23834.

55. Lee JB, Byeon JH, Jang HA, Kim JK, Yoo JW, Kikuchi Y, Lee BL. 2015. Bacterial cell motility of Burkholderia gut symbiont is required to colonize the insect gut. FEBS Lett 589:2784–2790.

56. Carlier A, Fehr L, Pinto-Carbó M, Schäberle T, Reher R, Dessein S, König G, Eberl L. 2016. The genome analysis of Candidatus Burkholderia crenata reveals that secondary metabolism may be a key function of the Ardisia crenata leaf nodule symbiosis. Environ Microbiol 18:2507–22.

57. Carlier AL, Eberl L. 2012. The eroded genome of a Psychotria leaf symbiont: hypotheses about lifestyle and interactions with its plant host. Environ Microbiol 14:2757–2769.

58. Christensen MJ, Bennett RJ, Ansari HA, Koga H, Johnson RD, Bryan GT, Simpson WR, Koolaard JP, Nickless EM, Voisey CR. 2008. Epichloë endophytes grow by intercalary hyphal extension in elongating grass leaves. Fungal Genet Biol 45:84–93.

59. Becker M, Becker Y, Green K, Scott B. 2016. The endophytic symbiont Epichloё festucae establishes an epiphyllous net on the surface of Lolium perenne leaves by development of an expressorium, an appressorium-like leaf exit structure. New Phytol 211:240–254.

60. Rutten PJ, Steel H, Hood GA, Ramachandran VK, McMurtry L, Geddes B, Papachristodoulou A, Poole PS. 2021. Multiple sensors provide spatiotemporal oxygen regulation of gene expression in a Rhizobium-legume symbiosis. PLoS Genet 17:e1009099.

61. Miyashiro T, Ruby EG. 2012. Shedding light on bioluminescence regulation in Vibrio fischeri. Mol Microbiol. NIH Public Access.

