## Supplementary information for "Distinct within-host bacterial populations ensure function, colonization and transmission in leaf symbiosis"

### Supplementary figures and tables

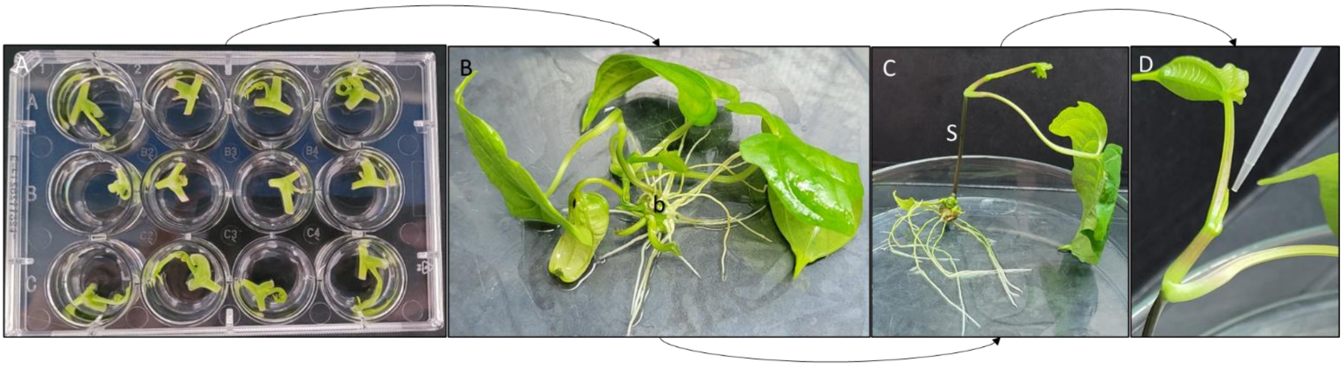


Figure S1: Method developed to make aposymbiotic plants and re-introduce a bacterium of interest

A: Node cuttings are taken from adult plants and incubated in a mixture of liquid MS, antibiotics and PPM for 3 weeks. B: After 3-4 weeks, a bulbil (b) with its root system become apparent. Multiple leaves have formed from the node and is providing sugars to the plant. C: The bulbil grows its own stem (S) that uses gravitropism to grow up and after the emergence of two leaves, the apical bud becomes visible. D: After confirmation of being aposymbiotic by crushing and plating out the newly developed acumen(s), the plant is re-inoculated with a bacterium of interest by dropping 2 µl of the bacterial suspension on the apical bud.

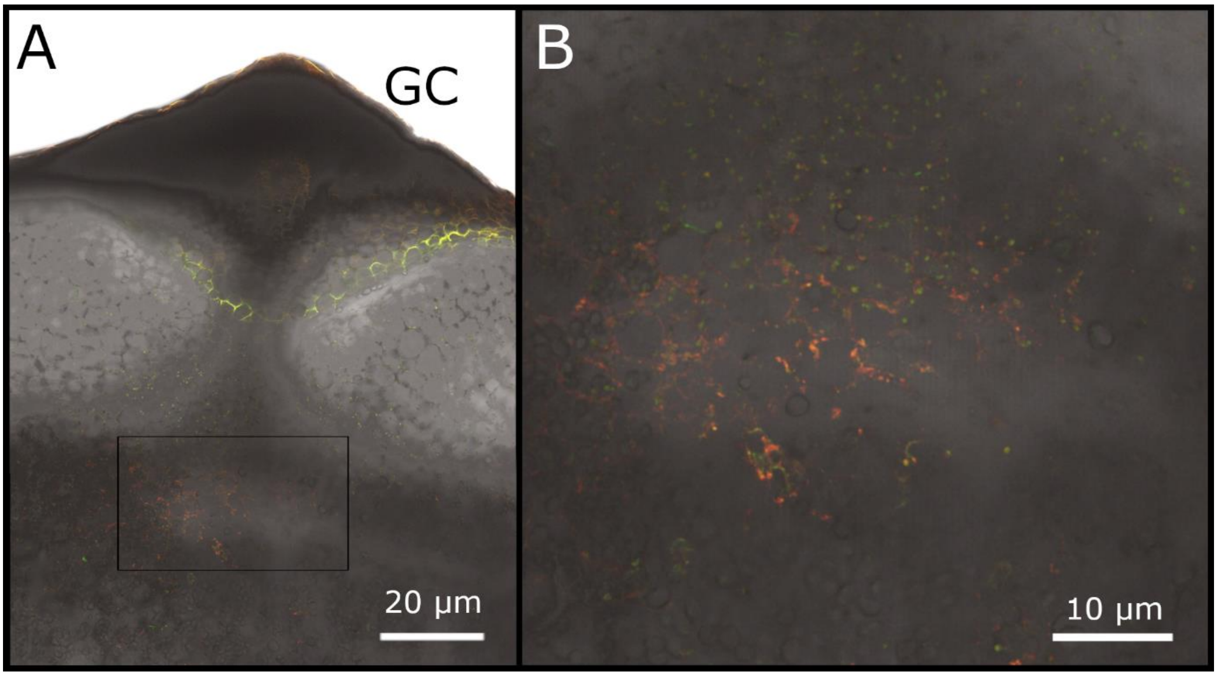


Figure S2: Colonization of mCherry-tagged *O. dioscoreae* in the bulbil.

Fresh section of growth primordium of *D. sansibarensis* bulbil, colonized by mCherry-tagged *O. dioscoreae,* imaged using confocal microscopy. Gnotobiotic plants were successfully colonized by mCherry-tagged *O. dioscoreae* and grown throughout the life cycle. Bulbils were harvested and after a few months, fresh section were cut through the ‘eyes’ of the bulbil. (A) The epidermis (ep) is loose from the underlying cell layers and a primordial plant structure emerges (hollow white arrow) from a pocket ± 1 cm from the surface. (B) Red fluorescence (arrows) can bee seen surrounding small plant cells, possibly meristematic tissue. Fluorescence seems to be diffused surrounding multiple cells, though concentrated in this one spot underneath the growth initial. Green autofluorescence can be seen.

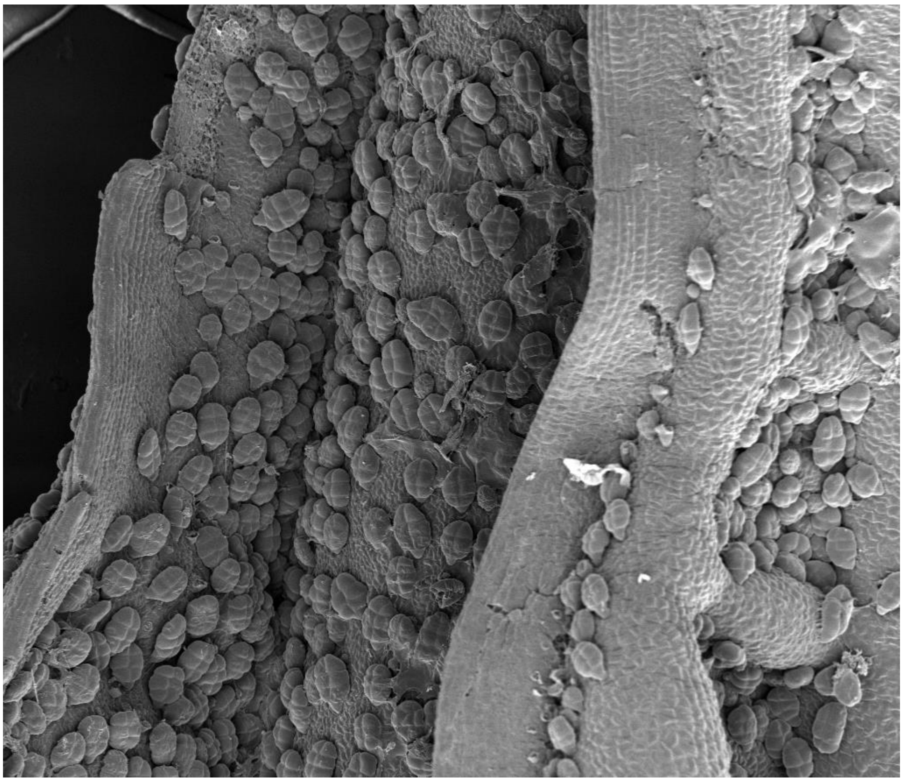

Figure S3: Adaxial leaf lamina of an older leaf primordium in the shoot tip, imaged by scanning electron microscopy.

Numerous glandular trichomes are present on the adaxial, and in lesser counts, on the abaxial side of the lamina. Glandular trichomes consist of 1 stalk cell and 5-6 glandular cells.

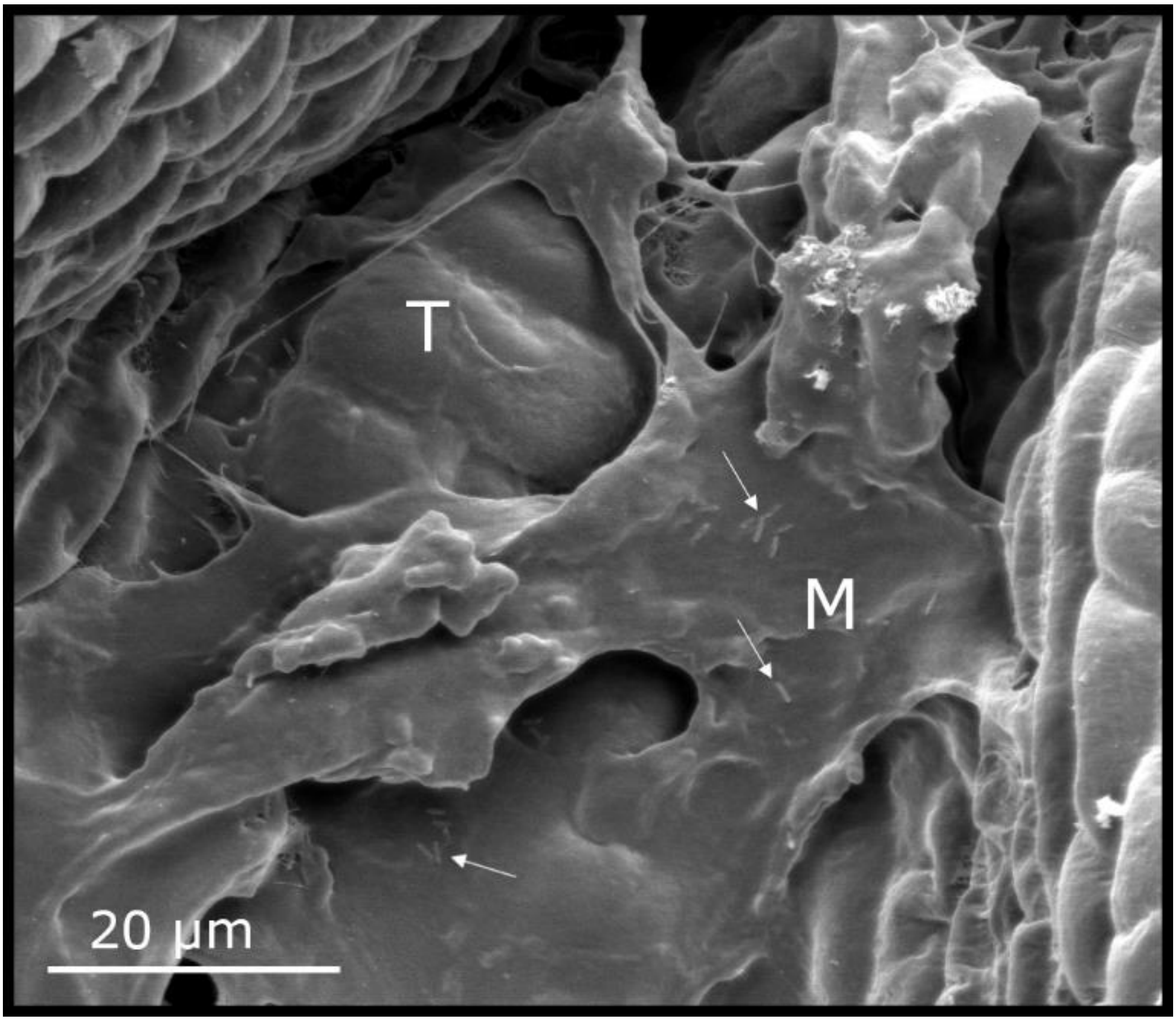


Figure S4: adaxial side of the leaf lamina of a primordial leaf in the shoot tip. Between the curling laminal sides, bacteria (arrows) reside in the mucus (M) that covers the trichomes (T).

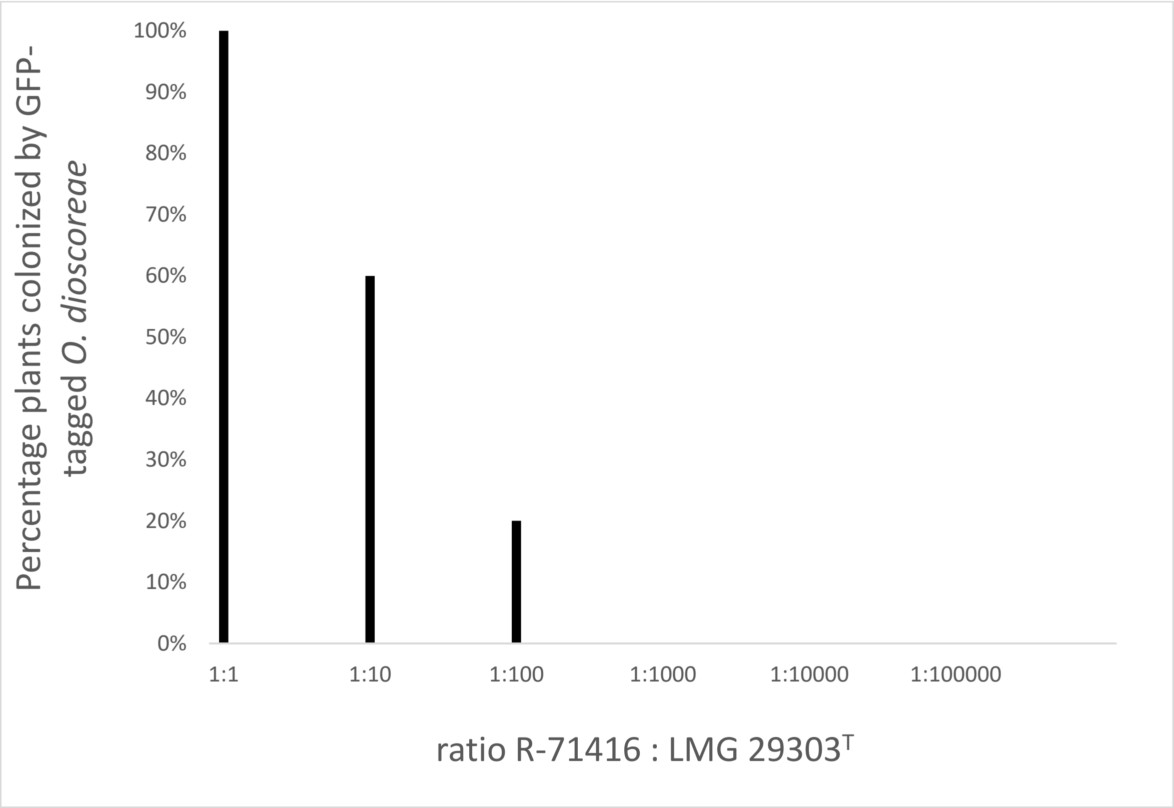


Figure S5: Percentage of *D. sansibarensis* plants colonized with GFP-tagged *O. dioscoreae* (strain R-71416) after co-inoculation with wild-type *O. dioscoreae* in diminishing ratios from 1:1 to 1:$10^5$.

*Table S1: Bacterial species and plasmids used in this study*

| **Species** | **Strain or plasmid** | **Description** | **Growth conditions** | **Reference or source** |
|---|---|---|---|---|
| | Strain | | | |
| *Orrella dioscoreae* | LMG 29303$^T$ | | TSA, 28°c, aerobic | (A. Carlier et al. 2017) |
| *Orrella dioscoreae* | R-71417 | Strain R71412 with mini Tn7(Gm)Ptac-mCherry | TSA + Nal30 + Gentamycin 20, 28°C, aerobic | This study |
| *Orrella dioscoreae* | R-71416 | Strain R71412 with mini Tn7(Gm)Ptac-GFP | TSA + Nal30 + Gentamycin 20, 28°C, aerobic | This study |
| *Orrella dioscoreae* | R-71412 | Spontaneous point mutation which made it resistant to Nalidixic acid | TSA + Nal30, 28°C, aerobic | (De Meyer et al., 2019) |
| *Escherichia coli* | mini-Tn7(Gm)Ptac-mCherry. | | LB broth+ Amp100 + Gm10, 37°C | (Choi & Schweizer, 2005) |
| *Escherichia coli* | mini Tn7(Gm)PA1/04/03-egfp-a | | LB broth+ Amp100 + Gm10, 37°C | (Choi & Schweizer, 2005) |
| *Orrella dioscoreae* | TA01 | MotB mutant: mCherry tagged Orrella with Kanamycin cassette in motB gene | TSA + Kanamycin 50 + Gentamycin 20 + Nalidixic acid 30, 28°C, aerobic | This study |
| *Orrella dioscoreae* | TA01 *motB*$^+$ | Strain TA01 harboring plasmid pBBR1MCS-3::*motAB* | TSA + Kanamycin 50 + Gentamycin 20 + Nalidixic acid 30, 28°C, aerobic | This study |
| *Orrella dioscoreae* | R-71416 | GFP-tagged Orrella dioscoreae with mini Tn7, derived from R-71412, Gentamycin resistant | TSA + Nal30 + Gentamycin 20, 28°C, aerobic | This study |

| | | | | |
|---|---|---|---|---|
| *Escherichia coli* | Top10 | Electrocompetent cells | LB broth, 37°C | |
| | Plasmids | | | |
| *Escherichia coli* | pDONRPEX18Tp-SceI-pheS | | LB broth + trimethoprim 25µg/ml, 37°C | (Fazli, Harrison, Gambino, Givskov, & Tolker-Nielsen, 2015) |
| *Escherichia coli* | pRK600 | pRK600 Helper plasmid in triparental conjugations; Cm-resistant ori-ColE1 RK-mob+ RK-tra | LB broth + Chloramphenicol 12.5 µg/ml, 37°C | (Kessler, de Lorenzo, & Timmis, 1992) |
| *Escherichia coli* | MT102 | pBBR1MCS-3 | | (Kovach et al., 1995) |
| *Escherichia coli* | Top10 | pBBR1MCS-3::*motAB* | LB broth, 37°C + tetracyclin 10µg/ml | This study |
| *Escherichia coli* | pUX-BF13 | *E. coli* SM10 lambda pir strain | LB broth + Ampiciline 100, 37°C | (Choi & Schweizer, 2005) |

*Table S2: Oligonucleotides*

| **Name** | **Sequence** |
|---|---|
| pA | AGAGTTTGATCCTGGCTCAG |
| pH | AAGGAGGTGATCCAGCCGCA |
| Mini Tn7 primer forward | GCC CTT TCG TCT TCA CCT CG |
| Mini Tn7 primer reverse | AGC TCC TGA AAA TCT CGC CA |
| GW-attB1 | GGGGACAAGTTTGTACAAAAAAGCAGGCT |
| GW-attB2 | GGGGACCACTTTGTACAAGAAAGCTGGGT |
| pKD4fwd-2 | TAGGCTGGAGCTGCTTCGAAGTTC |
| pKD4rev-2 | CATATGAATATCCTCCTTAGTTCCTATTCCG |
| motB-UpF-GW | TACAAAAAAGCAGGCTCTTCCTGGGCATTCTGCTT |
| motB-UpR-kan | GAACTTCGAAGCAGCTCCAGCCTAAGATCAGCCACAGCACGAG |

| | |
|---|---|
| motB-DnF-kan | CGGAATAGGAACTAAGGAGGATATTCATATGATGCGGTGCAGGAATTGAT |
| motB-DnR-GW | TACAAGAAAGCTGGGTCTTCCTGCAAGGTATCTGCATC |
| motAB-Fwd-KpnI | GGGGGTACCCATCTGTCGTCCGCCTCC |
| motAB-rev-SacI | GGGGAGCTCCATCCTCATCGTCACCGAA |
| nrdA-01-F | GAACTGGATTCCCGACCTGTTC |
| nrdA-02-R | TTCGATTTGACGTACAAGTTCTGG |
| gyrB-F | ACCAGCTTGTCCTTGGTCTG |
| gyrB-R | CGTGCTGTCGGTCAAGGT |
| nrp-F | AGGTATAGGGCACGATGAGC |
| nrp-R | CTGGATCTGCGCCACTTC |
| pqqc-F | AGGACTTCGGGCTGACACT |
| pqqc-R | TCGAACATGGTGTCGATGAG |
| KASII-F | GAGATCGCGGAAAACCAG |
| KASII-R | AAGAACAGGCCATCGACA |

*Table S3: Validation of differential regulation of select genes in shoot tip and acumen by quantitative RT-PCR. Ct values of housekeeping gene (gyrB 84-85, ODI_R4414) was used to normalise the Ct values of the operons of interest: smp1 (ODI_R1490), smp2 (ODI_R1505) and opk (ODI_R2249) using primers nrp 88-89, pqqc 90-91 and KASII 82-83, respectively. Ct values >40 indicate lack of detection after 40 PCR cycles. AB: Total RNA isolated from apical bud; LG: total RNA isolated from leaf gland.*

| **Ct** | **gyrB** | **smp1** | **smp2** | **opk** |
|---|---|---|---|---|
| **AB 1** | 33,91 | >40 | >40 | >40 |
| **AB 2** | 33,09 | >40 | >40 | >40 |
| **AB 3** | 33,12 | >40 | >40 | >40 |
| **AB 4** | 34,26 | 36,85 | 36,94 | >40 |
| **LG 1** | 32,55 | 32,78 | 30,46 | 33,49 |
| **LG 2** | 33,44 | 33,46 | 32,14 | 36,94 |
| **LG 3** | 32,99 | 31,26 | 29,34 | 33,87 |
| **LG 4** | 32,77 | 31,33 | 30,59 | 35,12 |
